# Integrative Analysis of Left Ventricle and Epicardial Adipose Tissue Identifies SDHA and OGDH as Candidate Targets for Ischemic Heart Disease

**DOI:** 10.1101/2025.09.17.676232

**Authors:** Muhammad Arif, Stephen Doran, Maryam Clausen, Johannes Wikström, Mohammad Bohlooly-Y, Elias Björnson, Liam Davidsson, Anders Jeppsson, Malin Levin, Adil Mardinoglu, Jan Boren

## Abstract

Ischemic heart disease (IHD) involves coordinated molecular changes across heart, yet their interplay remains poorly understood. Here, we investigated transcriptomic alterations in two heart tissue subtypes, left ventricle (LV) and epicardial adipose tissue (EAT), from age- and BMI-matched healthy and IHD individuals, including both diabetic and non-diabetic patients. We performed transcriptomic profiling and systems-level network analysis to identify disease-associated gene expression changes. Our analysis revealed: (1) stronger transcriptional responses in EAT compared to LV, particularly in diabetic individuals, and (2) widespread dysregulation of inflammatory and metabolic pathways, including oxidative phosphorylation, cytokine signaling, and fatty acid degradation, across both tissue subtypes. Co-expression network analysis uncovered shared gene modules, with SDHA (Succinate dehydrogenase complex, subunit A) and OGDH (Oxoglutarate Dehydrogenase) emerging as central, downregulated genes linked to mitochondrial function and inflammation, important processes in IHD pathophysiology. These findings were validated in independent human and mouse datasets. Overall, our integrative analysis identifies conserved molecular signatures across cardiac tissue subtypes, suggesting potential for therapeutic exploration in IHD.

**Graphical Abstract:** 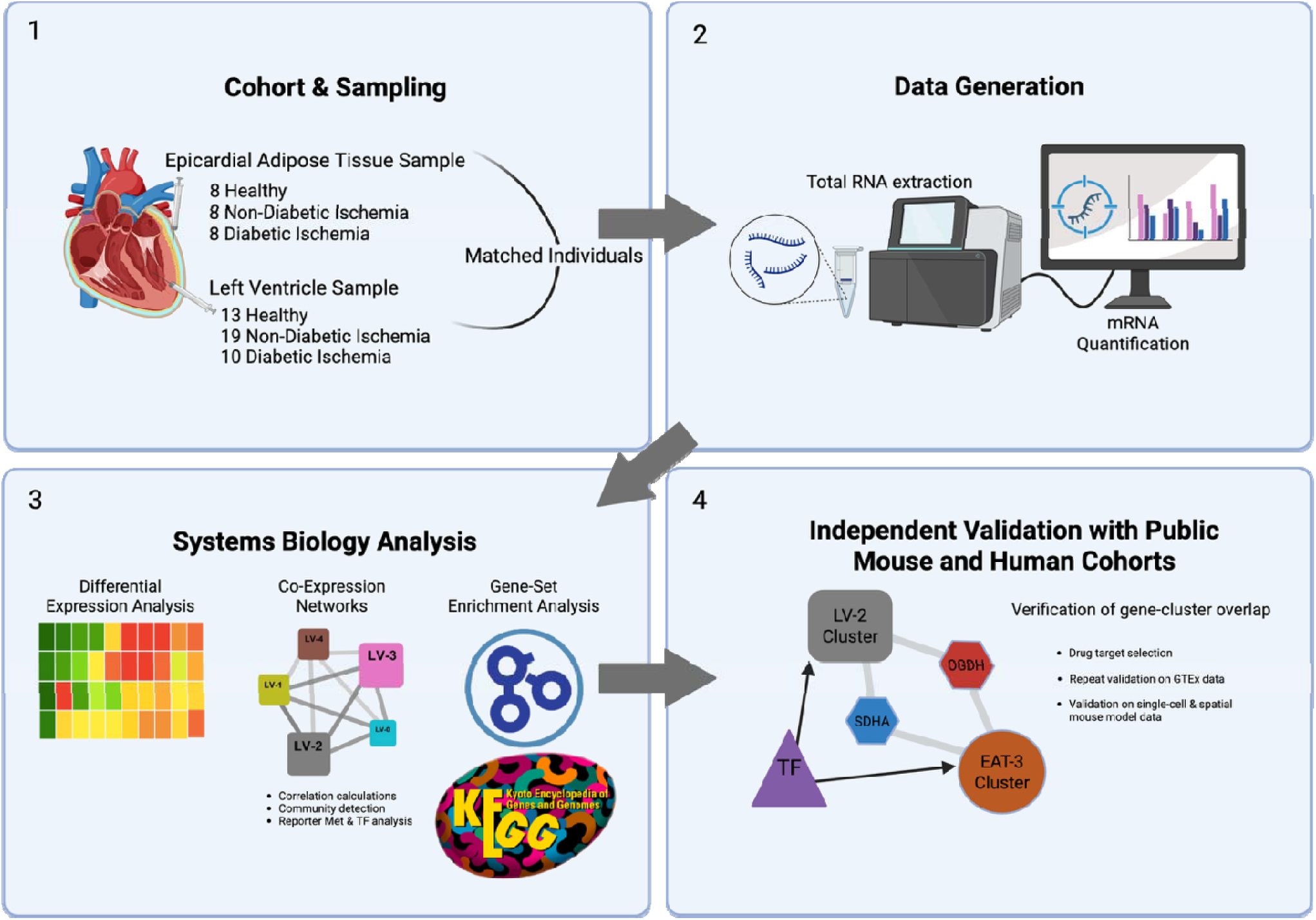

## Introduction

Ischemic heart disease (IHD) is a type of cardiovascular disorder caused by reduced blood and oxygen supply to the heart muscle. This reduction occurs due to the accumulation of plaque, particularly cholesterol deposits, within the walls of the coronary arteries, leading to their constriction. Based on data from WHO (WHO, 2025) and other international and national health bodies, IHD has been identified as the primary contributor to global mortality. In 2023, the Centers for Disease Control and Prevention (CDC) projected that 5% of adults (> 20 years old) have IHD in the United States, and 17% of IHD-related deaths happen in young and middle-aged adults (CDC, 2024). These numbers are likely to be underestimated as IHD is often silent and asymptomatic. A substantial portion of cases are only identified when patients experience a heart attack, a situation that frequently arises too late for effective intervention.

Despite extensive research on IHD, most transcriptomic studies have focused on a single tissue subtype such as atria or ventricles. Epicardial adipose tissue (EAT) is the visceral fat between the myocardium and the visceral pericardium and is linked to coronary events, notably in metabolic disease (Iacobellis, 2022; Jiang *et al*, 2017). EAT lacks a fascial barrier and shares microcirculation with the heart, enabling local paracrine effects on coronary arteries (Iacobellis, 2022). This anatomy distinguishes EAT from pericardial fat outside the parietal pericardium (Iacobellis, 2022). Higher EAT burden associates with incident coronary events independent of traditional risk factors (Mahabadi *et al*, 2013) and with faster coronary artery calcium (CAC) progression at follow-up (Mahabadi *et al*, 2014). Greater EAT volume also relates to high-risk plaque features on coronary CT angiography in coronary artery disease (CAD). Note that some imaging studies quantify pericoronary adipose tissue (PCAT), the perivascular subset of EAT. PCAT attenuation reflects local perivascular inflammation rather than total EAT burden (Iacobellis, 2022; Oikonomou *et al*, 2018). In cardiometabolic cohorts, EAT burden relates to CAD severity in type 2 diabetes independent of BMI and CAC, and EAT volume is higher in T2D with greater atherosclerosis (Mohar *et al*, 2014; Wang *et al*, 2009). Single-cell and single-nucleus studies of human EAT map inflammatory macrophage states along with fibro-adipogenic, endothelial and metabolic programs relevant to coronary pathology (Liu *et al*, 2024; Zou *et al*, 2023). Epidemiologic links between EAT burden and coronary events are observational, so a causal effect of lowering EAT on outcomes is unproven(Iacobellis, 2022; Mahabadi *et al*., 2013). These gaps make EAT a logical focus for cross-tissue signaling studies in IHD, but coordinated profiling with other cardiac compartments, such as left ventricle (LV), remains limited.

Mitochondrial dysfunction and metabolic remodelling are recognized among the hallmarks of IHD and heart failure (Bertero & Maack, 2018; Brown *et al*, 2017; Liu *et al*, 2022; Zhou & Tian, 2018). Among the central players in these processes, succinate dehydrogenase complex flavoprotein subunit A (SDHA) and oxoglutarate dehydrogenase (OGDH) have been implicated in heart diseases. SDHA mutation has been associated to genetic heart diseases(Courage *et al*, 2017; Levitas *et al*, 2010), while cardiac OGDH level has been linked to hypertrophic cardiomyopathy severity and proposed as a potential atherosclerosis biomarker(Moskowitz, 1986; Ranjbarvaziri *et al*, 2021). These prior findings underscore the importance of mitochondrial metabolism in IHD and support the rationale for investigating SDHA and OGDH as potential therapeutic targets.

In this study, we performed transcriptomics analysis of two distinct cardiac tissue subtypes, the LV and EAT from healthy individuals and those with ischemic heart disease (IHD) (**Figure S1**). We hypothesized that IHD induces tissue subtype- and context-specific transcriptional and network alterations, particularly in mitochondrial metabolic pathways, and that these changes are further modulated by diabetes. By comparing diabetic IHD, non-diabetic IHD, and healthy control subjects, and by analyzing LV and EAT from the same hearts, we uniquely evaluated shared and distinct molecular networks across regions and disease states to identify key drivers of IHD pathophysiology. Using systems biology and co-expression network analysis, we uncovered interactions between LV and EAT, common gene clusters, and central genes linked to disease pathology. Notably, while SDHA and OGDH have previously been implicated in cardiac mitochondrial dysfunction and metabolic remodelling, our study demonstrates their consistent emergence as central nodes across LV and EAT, in both diabetic and non-diabetic IHD. This highlights their potential as robust candidate biomarkers and therapeutic targets within regulatory networks of IHD.

## Results

### Characteristics of the study participants

This study aimed to identify transcriptional differences in the heart between healthy control and subjects with ischemic heart diseases, in both diabetic and non-diabetic subjects. We recruited 14 healthy control subjects and 30 subjects with ischemic heart diseases (IHD) (**Data S1**). One control subject was excluded due to extremely high glucose level and one IHD patient was removed due to inconsistency in the sequencing data. The remaining 29 IHD patients were then split into two subgroups, the non-diabetic (ND-I, n = 19) and diabetic (D-I, n = 10) groups. We observed no significant differences in the groups’ age, body mass index, and other clinical variables, except plasma glucose, systolic blood pressure, and serum potassium level (**Table 1, Data S1**). As expected, the glucose level of the diabetic patients was the highest compared to other groups, together with their systolic blood pressure (**Table 1**).

**Table 1.**
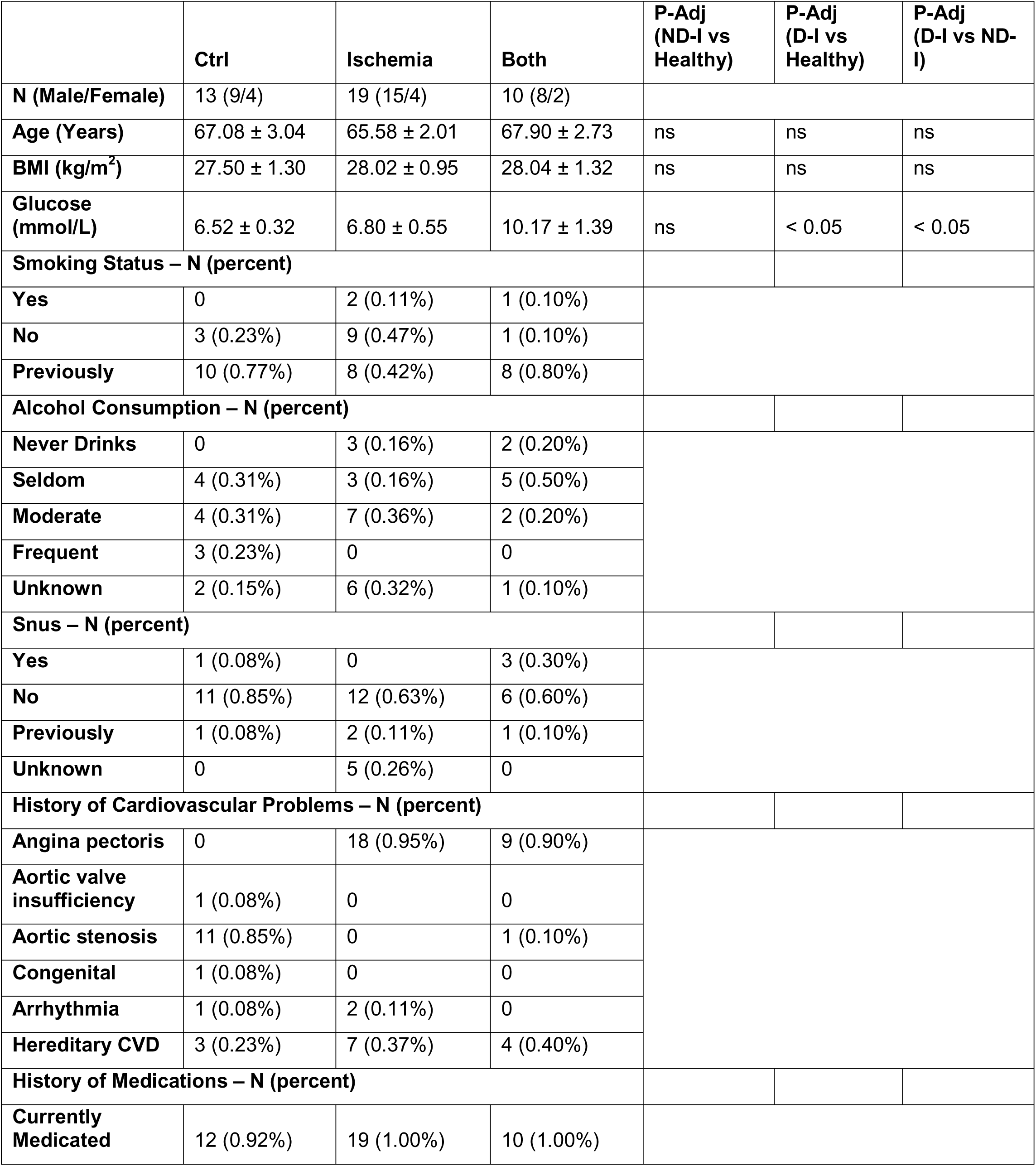
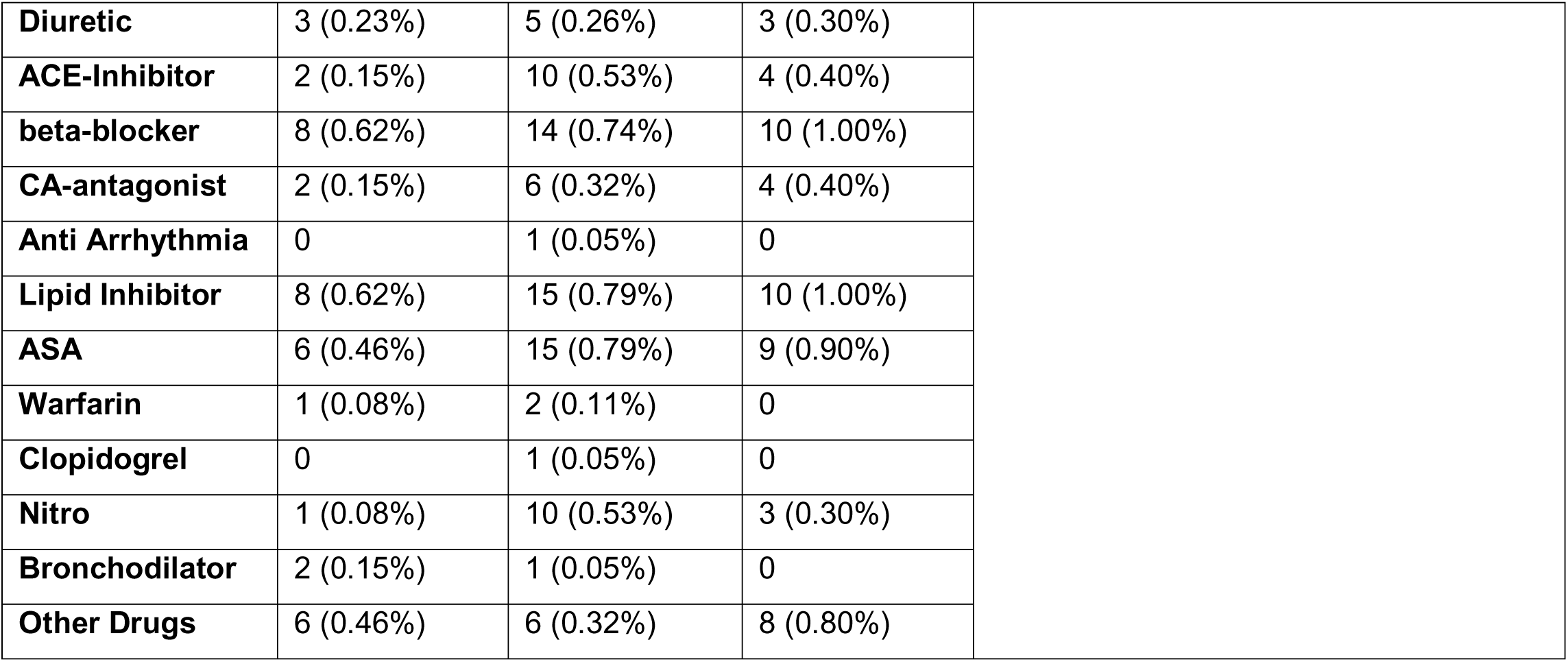
Demographic and clinical characteristics of the participants.

|  | Ctrl | Ischemia | Both | P-Adj<br>(ND-I vs<br>Healthy) | P-Adj<br>(D-I vs<br>Healthy) | P-Adj<br>(D-I vs ND-I) |
| --- | --- | --- | --- | --- | --- | --- |
| **N (Male/Female)** | 13 (9/4) | 19 (15/4) | 10 (8/2) |  |  |  |
| **Age (Years)** | 67.08 ± 3.04 | 65.58 ± 2.01 | 67.90 ± 2.73 | ns | ns | ns |
| **BMI (kg/m<sup>2</sup>)** | 27.50 ± 1.30 | 28.02 ± 0.95 | 28.04 ± 1.32 | ns | ns | ns |
| **Glucose**<br>**(mmol/L)** | 6.52 ± 0.32 | 6.80 ± 0.55 | 10.17 ± 1.39 | ns | < 0.05 | < 0.05 |
| **Smoking Status – N (percent)** |  |  |  |  |  |  |
| **Yes** | 0 | 2 (0.11%) | 1 (0.10%) |  |  |  |
| **No** | 3 (0.23%) | 9 (0.47%) | 1 (0.10%) |  |  |  |
| **Previously** | 10 (0.77%) | 8 (0.42%) | 8 (0.80%) |  |  |  |
| **Alcohol Consumption – N (percent)** |  |  |  |  |  |  |
| **Never Drinks** | 0 | 3 (0.16%) | 2 (0.20%) |  |  |  |
| **Seldom** | 4 (0.31%) | 3 (0.16%) | 5 (0.50%) |  |  |  |
| **Moderate** | 4 (0.31%) | 7 (0.36%) | 2 (0.20%) |  |  |  |
| **Frequent** | 3 (0.23%) | 0 | 0 |  |  |  |
| **Unknown** | 2 (0.15%) | 6 (0.32%) | 1 (0.10%) |  |  |  |
| **Snus – N (percent)** |  |  |  |  |  |  |
| **Yes** | 1 (0.08%) | 0 | 3 (0.30%) |  |  |  |
| **No** | 11 (0.85%) | 12 (0.63%) | 6 (0.60%) |  |  |  |
| **Previously** | 1 (0.08%) | 2 (0.11%) | 1 (0.10%) |  |  |  |
| **Unknown** | 0 | 5 (0.26%) | 0 |  |  |  |
| **History of Cardiovascular Problems – N (percent)** |  |  |  |  |  |  |
| **Angina pectoris** | 0 | 18 (0.95%) | 9 (0.90%) |  |  |  |
| **Aortic valve**<br>**insufficiency** | 1 (0.08%) | 0 | 0 |  |  |  |
| **Aortic stenosis** | 11 (0.85%) | 0 | 1 (0.10%) |  |  |  |
| **Congenital** | 1 (0.08%) | 0 | 0 |  |  |  |
| **Arrhythmia** | 1 (0.08%) | 2 (0.11%) | 0 |  |  |  |
| **Hereditary CVD** | 3 (0.23%) | 7 (0.37%) | 4 (0.40%) |  |  |  |
| **History of Medications – N (percent)** |  |  |  |  |  |  |
| **Currently**<br>**Medicated** | 12 (0.92%) | 19 (1.00%) | 10 (1.00%) |  |  |  |

|  |  |  |  |
| --- | --- | --- | --- |
| **Diuretic** | 3 (0.23%) | 5 (0.26%) | 3 (0.30%) |
| **ACE-Inhibitor** | 2 (0.15%) | 10 (0.53%) | 4 (0.40%) |
| **beta-blocker** | 8 (0.62%) | 14 (0.74%) | 10 (1.00%) |
| **CA-antagonist** | 2 (0.15%) | 6 (0.32%) | 4 (0.40%) |
| **Anti Arrhythmia** | 0 | 1 (0.05%) | 0 |
| **Lipid Inhibitor** | 8 (0.62%) | 15 (0.79%) | 10 (1.00%) |
| **ASA** | 6 (0.46%) | 15 (0.79%) | 9 (0.90%) |
| **Warfarin** | 1 (0.08%) | 2 (0.11%) | 0 |
| **Clopidogrel** | 0 | 1 (0.05%) | 0 |
| **Nitro** | 1 (0.08%) | 10 (0.53%) | 3 (0.30%) |
| **Bronchodilator** | 2 (0.15%) | 1 (0.05%) | 0 |
| **Other Drugs** | 6 (0.46%) | 6 (0.32%) | 8 (0.80%) |

From the remaining 42 subjects (13 healthy control and 29 IHD), we collected left ventricular (LV) tissue. In addition, from a subset of 24 subjects (n = 8 per group), we also collected matched epicardial adipose tissue (EAT). In total, this resulted in 66 transcriptomic samples (42 LV and 24 matched EAT). Transcriptomic profiling was performed on these samples (**Data S1**).

### Left ventricle of ischemic subjects shows a significant increase in inflammatory responses and a decrease in carbohydrate metabolism pathways

First, we performed a differential expression analysis of the LV transcriptomics data (**Data S2**). We identified 839 (436 up and 403 down-regulated) and 755 (368 up and 387 down-regulated) differentially expressed genes (DEGs) when comparing ND-I and D-I groups to healthy subjects, respectively (**Figure 1A**). To understand the affected biological functions associated with these transcriptomics changes, we performed gene-set enrichment analysis (GSEA) to identify the association of the DEGs with biological pathways and processes (**Data S2**). The GSEA using KEGG pathways in ND-I subjects showed up-regulation of sulfur metabolism, spliceosome, and mineral absorption pathways compared to the healthy group (**Figure 1B**). On the other hand, we observed significant down-regulation in several metabolic pathways in ND-I samples compared to healthy subjects, specifically lysine degradation, arginine and proline metabolism, insulin signaling, glycolysis/glucogenesis, and TCA cycle. We observed bigger changes in the pathways when comparing D-I samples with healthy control (**Figure 1C**). Specifically, inflammatory and cell death pathways were up-regulated, e.g. NF-kappa B signaling, natural killer cell signaling, Fc gamma R-mediated phagocytosis, B and T cell receptor signaling pathways, and cellular senescence, while oxidative phosphorylation pathway was down-regulated. We also observed the up-regulation of pathways associated with lipid metabolism (PPAR signaling, sphingolipid biosynthesis, and steroid pathways) and fatty acid degradation, and the down-regulation of glycine, serine, and threonine metabolisms. Also, we observed similarities in the alteration caused by both conditions. Both conditions showed down-regulation of carbohydrate metabolism pathways and up-regulation of important inflammation and immune response pathways, specifically TNF, Chemokine, JAK-STAT, Cytokine-cytokine receptor, and Toll-like receptor signaling pathways (**Figure 1C**). We further confirmed these alterations by performing GSEA on Gene Ontology biological process and observed similar changes in the associated terms **(Figure S1**: Graphical Abstract and Study Flow Figure S2).

**Figure 1:**
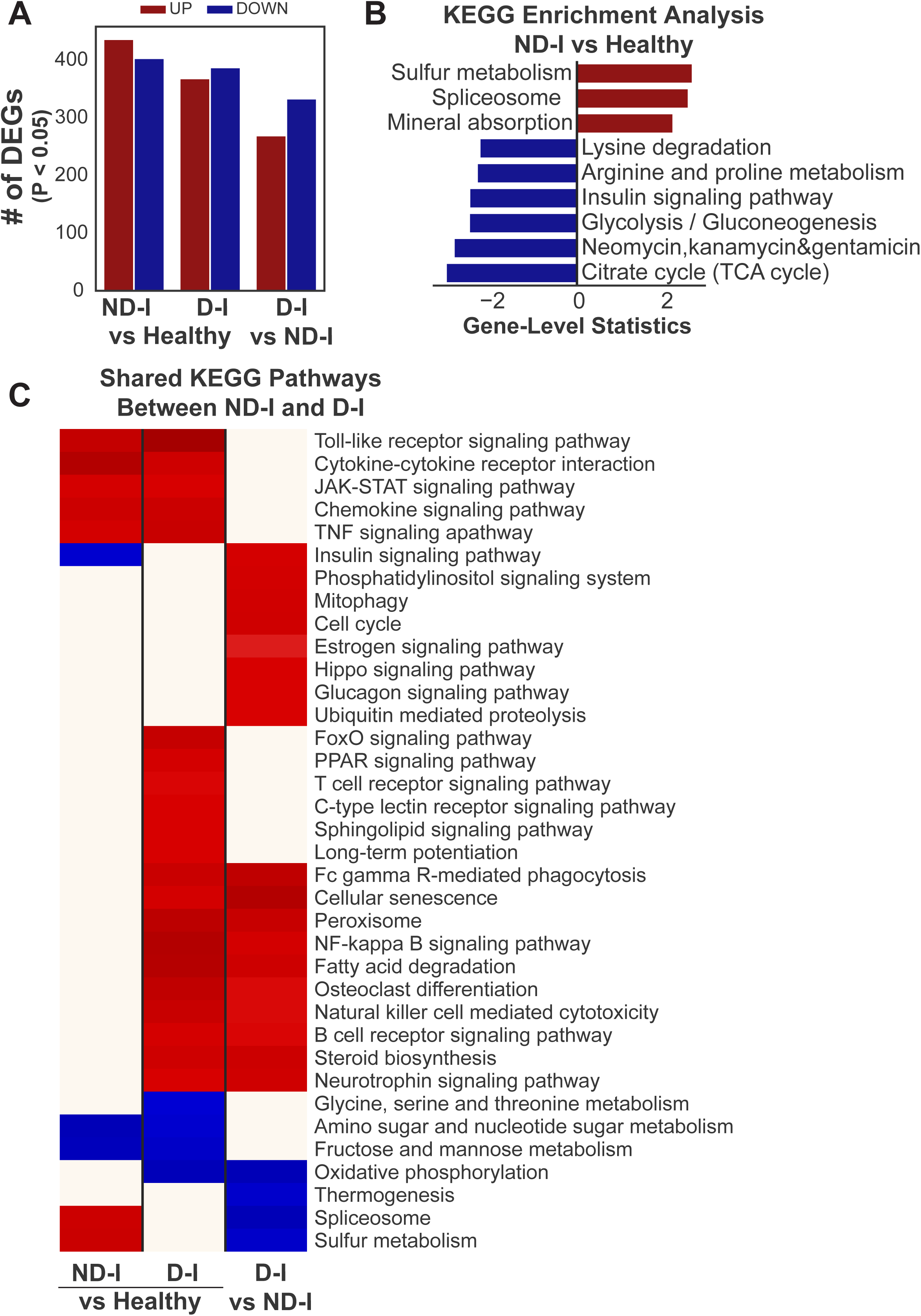
Transcriptomics of Heart Left Ventricle. **(A)** The differential expression analysis showed alterations in significant transcriptional profile alterations (P-Value < 0.05) between the ischemic (ND-I and D-I) and healthy subjects. **(B)** KEGG pathways (P-Value < 0.05) that were uniquely altered when comparing ND-I with healthy subjects. **(C)** Shared KEGG pathways (FDR < 0.01) were significantly altered when comparing D-I subjects with healthy and ND-I subjects.

We also compared the D-I subjects with ND-I. There were 269 up and 333 down-regulated genes in this comparison (**Figure 1A**). Although many of the altered pathways were similar to D-I vs healthy comparisons, we observed that several pathways were uniquely altered in the D-I compared to ND-I subjects (**Figure 1C**), including the up-regulation of phosphatidylinositol, estrogen, hippo, and glucagon signaling pathways, and cellular processes (mitophagy and cell cycle), while thermogenesis was down-regulated.

### Ischemic epicardial adipose tissue exhibits more prominent transcriptional alterations compared to the left ventricle

We performed a similar analysis for the EAT transcriptomics data **(Data S3)** and, interestingly, observed significantly higher DEGs compared to LV, especially in D-I subjects. We identified 1109 (545 up and 564 down-regulated) and 1317 (801 up and 516 down-regulated) DEGs in ND-I and D-I subjects when compared to the healthy group (**Figure 2A**). Functionally, EAT showed a more pronounced effect in their response to IHD (**Figure 2B, Data S3**). In ND-I subjects, we observed more down-regulated metabolic pathways associated with lipids, amino acids, carbohydrates, and other important signaling pathways. The altered metabolic pathways in LV were also found to be down-regulated in EAT (**Figure 2B**). The D-I subjects showed similar, but more pronounced, up-regulation of inflammatory and cell death pathways as LV (**Figure 2B**). Moreover, we observed a fibrosis-related pathway, cell adhesion molecule, to be up-regulated in D-I subjects compared to healthy, while the butanoate metabolism pathway was down-regulated (**Figure 2B**).

**Figure 2:**
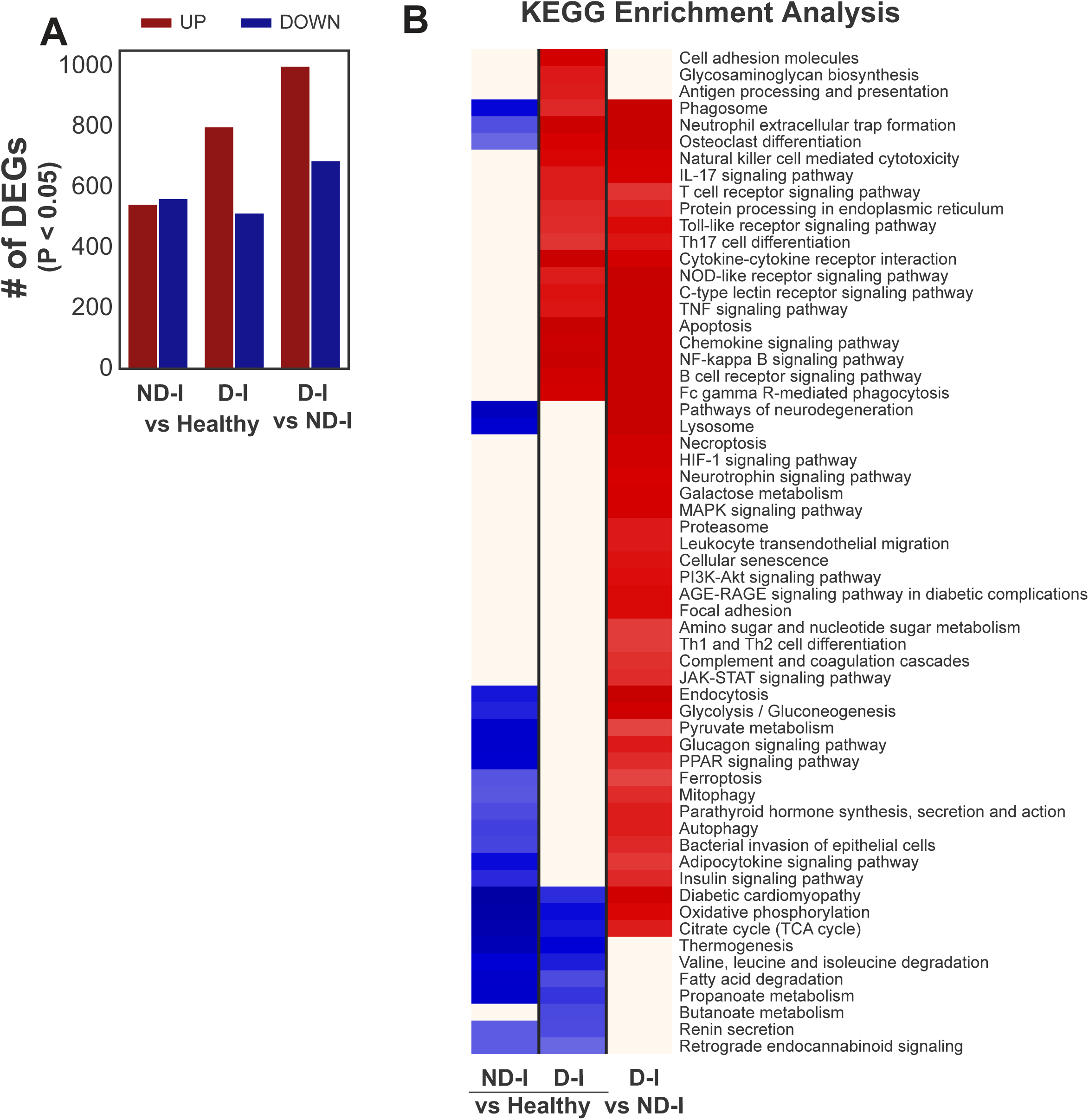
Transcriptomics of Epicardial Adipose Tissue. **(A)** The differential expression analysis showed alterations in significant transcriptional profile alterations (P-Value < 0.05) between the ischemic (ND-I and D-I) and healthy subjects. **(B)** KEGG pathways (FDR < 0.01) that are uniquely altered when comparing ND-I and D-I with healthy subjects, and ND-I vs D-I.

We also identified multiple commonly regulated pathways in both ND-I and D-I groups when compared to healthy subjects (**Figure 2B, Figure S3**). Mitochondrial functions and oxidative phosphorylation were down-regulated, together with diabetic cardiomyopathy and thermogenesis pathways. Moreover, we identified several metabolic pathways to be commonly down-regulated in both conditions, such as the TCA cycle, valine, leucine, and isoleucine degradation, fatty acid degradation, propanoate metabolism, and retrograde endocannabinoid signaling pathways. Interestingly, we observed that many known fibroproliferative pathways and processes, such as extracellular matrix organization and osteoclast differentiation, were up-regulated in the D-I group but down-regulated in the ND-I group, together with antigen processing and presentation and neutrophil-related immune response (**Figure 2B, Figure S3**).

Furthermore, we compared D-I to ND-I subjects and observed high dysregulation in transcriptomics data of EAT (1001 up and 689 down-regulated genes) (**Figure 2A**). Functional analyses showed similar dysregulation as D-I and healthy comparison (**Figure 2B**), several pathways were uniquely significantly up-regulated in D-I compared to ND-I subjects, including signaling pathways (HIF-1, MAPK, PI3K-Akt, and JAK-STAT), immune (Th1 and Th2 cell differentiation and leukocyte transendothelial migration), focal adhesion, and cellular senescence. This showed that, although both have similar disease etiology, D-I subjects have higher biological alterations compared to ND-I subjects.

### Gene co-expression network unveils tissue subtype-specific clusters associated with ischemic heart diseases

In the previous sections, we successfully identified the transcriptional alterations and affected biological pathways in response to ischemic heart diseases in both LV and EAT. To further study the molecular functional relationships and identify driver pathways associated with the disease, we employed co-expression networks (CNs). CNs are a powerful tool in untangling the complexity of big biological data in healthy and diseases (Arif *et al*, 2021b), including cardiovascular (Arif *et al*, 2021a), fibrotic (Arif *et al*, 2023), and metabolic diseases (Zeybel *et al*, 2022). Here, we generated two tissue subtype-specific CNs (LV and EAT) and performed downstream network analysis on the top 25% significantly positively co-expressed genes (FDR < 0.05) in each network. We employed the Leiden community detection algorithm (Traag *et al*, 2019) to better understand the network by identifying clusters with distinct network structures. Furthermore, we calculated the average transitivity of the clusters to identify the most central clusters in each network (Lee *et al*, 2017). We hypothesized that the most central clusters play important roles in responding to the disease, thus their gene members are rational therapeutic target candidates.

Using the LV transcriptomics data, we generated an LV-specific CN with 13733 gene nodes connected with 1.6M undirected edges. From the CN, we identified 5 distinct clusters (**Figure 3A**) with the biggest cluster, LV-0, containing 5284 genes and the smallest, LV-4, with 439 genes. Interestingly, based on their average transitivity (**Data S4**), we found LV-2 (3073 genes) and LV-3 (1787) as the central and driver clusters of the LV-specific CN. Similarly, we generated EAT-specific CN. It has 12934 gene nodes connected with more than 258K undirected edges and identified 9 distinct clusters (**Figure 3B**) with EAT-0 and EAT-8 as the biggest and smallest clusters, with 3821 and 59 genes, respectively. EAT-3 and EAT-5 showed the highest average transitivity; thus, we defined them as the most central and driver network in EAT-specific CN.

**Figure 3:**
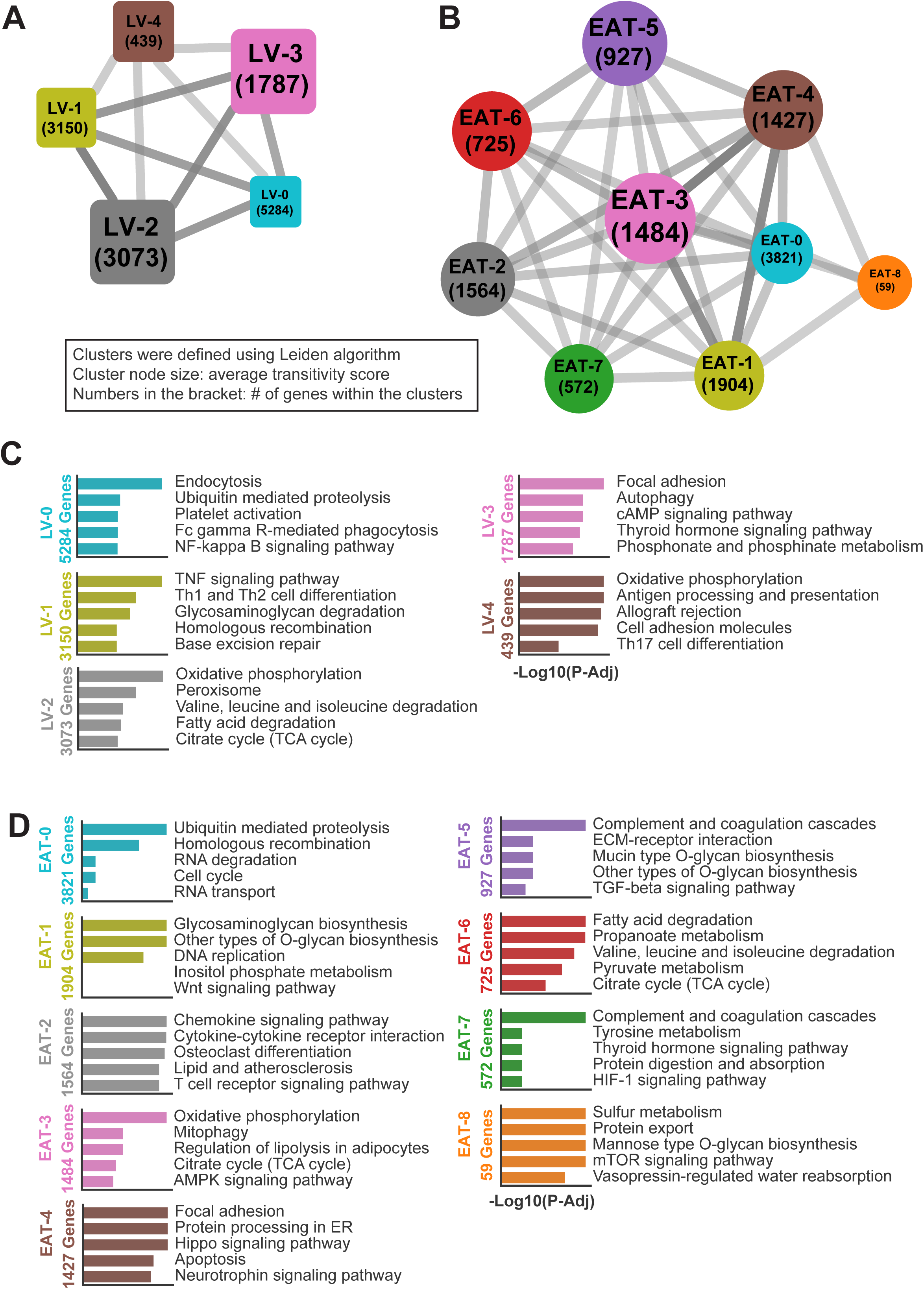
Heart tissue subtype-specific co-expression networks (CNs) **(A)** LV-specific CN consisted of five clusters with distinct gene members with LV-2 and LV-3 as the two most central clusters, based on their average transitivity scores (node size). The numbers below the name denote the number of genes within the clusters. **(B)** EAT-specific CN consisted of 9 clusters with EAT-3 and EAT-5 as the two most central clusters **(C)** Top 5 associated KEGG pathways of each cluster of LV-specific CN. **(D)** Top 5 associated KEGG pathways of each cluster of EAT-specific CN.

To further understand the cluster roles in the disease, we performed GSEA with KEGG Pathways on each cluster to identify the functions associated with their gene members (**Figure 3C-D**). Interestingly, the central clusters from both tissue subtype-specific CNs also showed top similar functional roles. LV-2 and EAT-3 were enriched with genes associated with oxidative phosphorylation. Moreover, the genes from both clusters were also associated with an important metabolic pathway, the TCA cycle, and pathways associated with lipid metabolism, specifically peroxisome and fatty acid degradation in LV-2 and regulation of lipolysis in adipocytes in EAT-3. Uniquely, LV-2 was enriched with genes associated with valine, leucine, and isoleucine degradation (**Figure 3C**) and EAT-3 with mitophagy and AMPK signaling pathways (**Figure 3D**). The other central clusters, LV-3 and EAT-5 were enriched with fibrosis-related pathways: focal adhesion and ECM-receptor interaction (**Figure 3C-D**). LV-3 was also enriched with genes related to autophagy, cAMP signaling, thyroid hormone signaling, and phosphonate and phosphinate metabolism pathways (**Figure 3C**), and EAT-3 with complement and coagulation cascades, glycan biosynthesis, and TGF-beta signaling pathway (**Figure 3D**).

### Reporter metabolite analysis of CN connects central clusters to NAD metabolism and ubiquinone and ubiquinol metabolites

Since the functional analyses from differential expression and network analyses showed a high association between ischemic heart disease and metabolic pathways, we performed reporter metabolite analysis on each cluster to understand the cluster gene members association with metabolites. We found 828 metabolites significantly associated with LV-0, LV-2, LV-4, EAT-0, EAT-3, EAT-6, and EAT-7 (**Figure S4**). We further isolated the metabolites that were associated with at least 2 clusters to examine the clusters’ commonalities (**Figure 4**). We focused on LV-2 and EAT-3, as these were the only central clusters with significantly associated reporter metabolites. By far, LV-2 showed the highest connectivity with 819 metabolites and shared them with all other clusters (**Figure S4**). On the other hand, EAT-3 is only connected to 8 metabolites (**Figure S4**), and all of them are also connected to LV-2 (**Figure 4**), including important metabolites such as NAD, NADH, and FAD. Moreover, they were also connected to sulfide and iron, known central metabolites of sulfur metabolism and mineral absorption, respectively. Finally, we observed that gene members of LV-2 and EAT-3 were also associated with Coenzyme Q, Ubiquinone Q1, and QH2 (ubiquinol) (**Figure 4A**).

**Figure 4:**
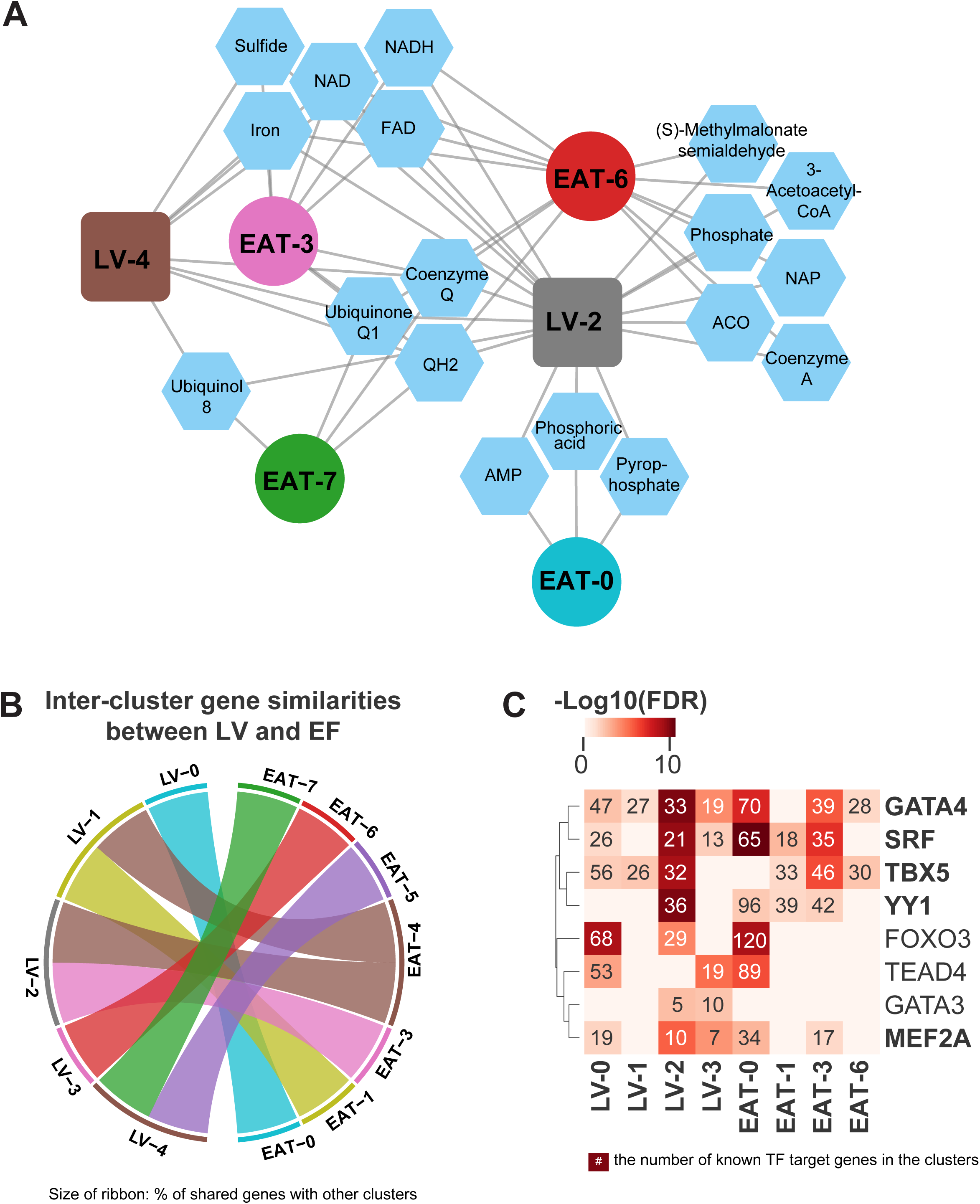
Reporter Metabolites and Commonalities between LV and EAT-specific CN clusters. **(A)** Reporter metabolites analysis results connected to their associated CN clusters. Only metabolites and clusters with degree centrality > 1 were shown. For a more complete figure, refer to **Figure S4 (B)** Hypergeometric test results revealed the inter-cluster gene similarities between LV and EAT-specific CNs. **(C)** Known cardiac transcription factors and their regulated clusters in both regions CNs. The annotations denoted the number of known TF target genes from the clusters. Bold gene names denoted the TFs regulating both LV-2 and EAT-3.

### LV-2 and EAT-3 are regulated by known cardiac transcription factors and exhibit potential as the location of therapeutic targets

Based on the functional and reporter metabolite similarities of the central clusters described above, we assumed that the CN of these two distinct heart tissue subtypes was driven by similar genes. To validate that, we performed the hypergeometric test to identify these similarities (**Data S4**). We found that indeed LV-2 and EAT-3 shared a significant number of genes (**Figure 4B**) but not between LV-3 and EAT-5. Based on the facts that LV-2 and EAT-3 were both identified as central clusters and shared important genes and pathways related to the diseases, we hypothesized that one or more shared genes in LV-2 and EAT-3 might be the rational therapeutic targets to treat ischemic heart diseases, for both diabetic and non-diabetic patients.

To identify candidate therapeutic targets, first, we were interested in the common transcription factors (TFs) that regulate LV-2 and EAT-3. We performed reporter TFs analysis (Huang *et al*, 2017) on all clusters (**Data S4**) and successfully mapped 170 TFs to the 14 LV and EAT clusters (**Data S4**). We identified 68 TFs that are commonly associated with LV-2 and EAT-3 clusters (**Figure S5**). Interestingly, out of the 8 known cardiac TFs that we successfully mapped to the clusters, 5 of them were mapped, among others, to both LV-2 and EAT-3, namely GATA4, SRF, TBX5, YY1, and MEAT2A (**Figure 4C**).

### Identification of candidate therapeutic targets for ischemic heart diseases

Based on the previous analyses, although we had two distinct tissue subtypes of heart, we successfully showed the similarities in their response to ischemic heart disease. Moreover, these analyses supported the hypothesis that gene members of the central clusters, specifically LV-2 and EAT-3, may represent potential targets for mitigating the effects of ischemic heart disease. To pinpoint the target candidates, we narrowed down the common genes in LV-2 and EAT-3 and examined if they are associated with the shared reporter metabolites or known cardiac TFs, mentioned in the previous sections, and we found 20 genes (**Data S5**) that fulfilled all those criteria. To facilitate pre-clinical research, we examined the expression changes of these genes in three mouse myocardial infarction publicly available data (Arif *et al*., 2021a; Ounzain *et al*, 2015; Williams *et al*, 2018) and found 4 genes to be differentially expressed in all cohorts:

SDHA, OGDH, SDHC, and FLAD1. We decided to focus on SDHA (Succinate dehydrogenase complex flavoprotein subunit A) and OGDH (Oxoglutarate dehydrogenase) as they are also FDA-approved drug targets, based on the information from the Human Protein Atlas (https://www.proteinatlas.org/)(Uhlen *et al*, 2015). SDHA and OGDH were both the target of two main known cardiac TFs, SRF and TBX5. Furthermore, OGDH was associated with two reporter metabolites, NAD and NADH, while SDHA is known to be associated with the rest of the reporter metabolites.

To further understand their functional roles, we isolated SDHA, OGDH, and their top 10 neighbors from LV and EAT-specific networks and their shared neighbors (**Figure 5A**) that were significantly altered in ND-I or D-I compared to healthy patients. In LV, we observed that SDHA and OGDH were a direct neighbor (high correlation) and shared 11 neighboring genes that were down-regulated in ND-I or D-I, including AIFM1 (associated with apoptosis), PCYT2 and LMF1 (lipid metabolism), TIMM23B and TIMM44 (mitochondria), PSMB5 (proteasome), ENGASE and SLC2A11 (carbohydrate metabolism), and NDUFS2 (mitochondria, NADH, and FAD activity, also diabetic cardiomyopathy). Interestingly, when we checked their shared neighbors in EAT, we found a unique set of 11 down-regulated genes, including MCAM (vascular wound healing), RAB1B (glycoprotein metabolism), NDUFV1 (mitochondria), and ARHGEF7 (known to be related to cardiac arrest and coronary heart disease). The functional relevance of SDHA’s and OGDH’s neighbors to cardiovascular diseases further reinstate their potential role in ischemic heart disease pathology.

**Figure 5:**
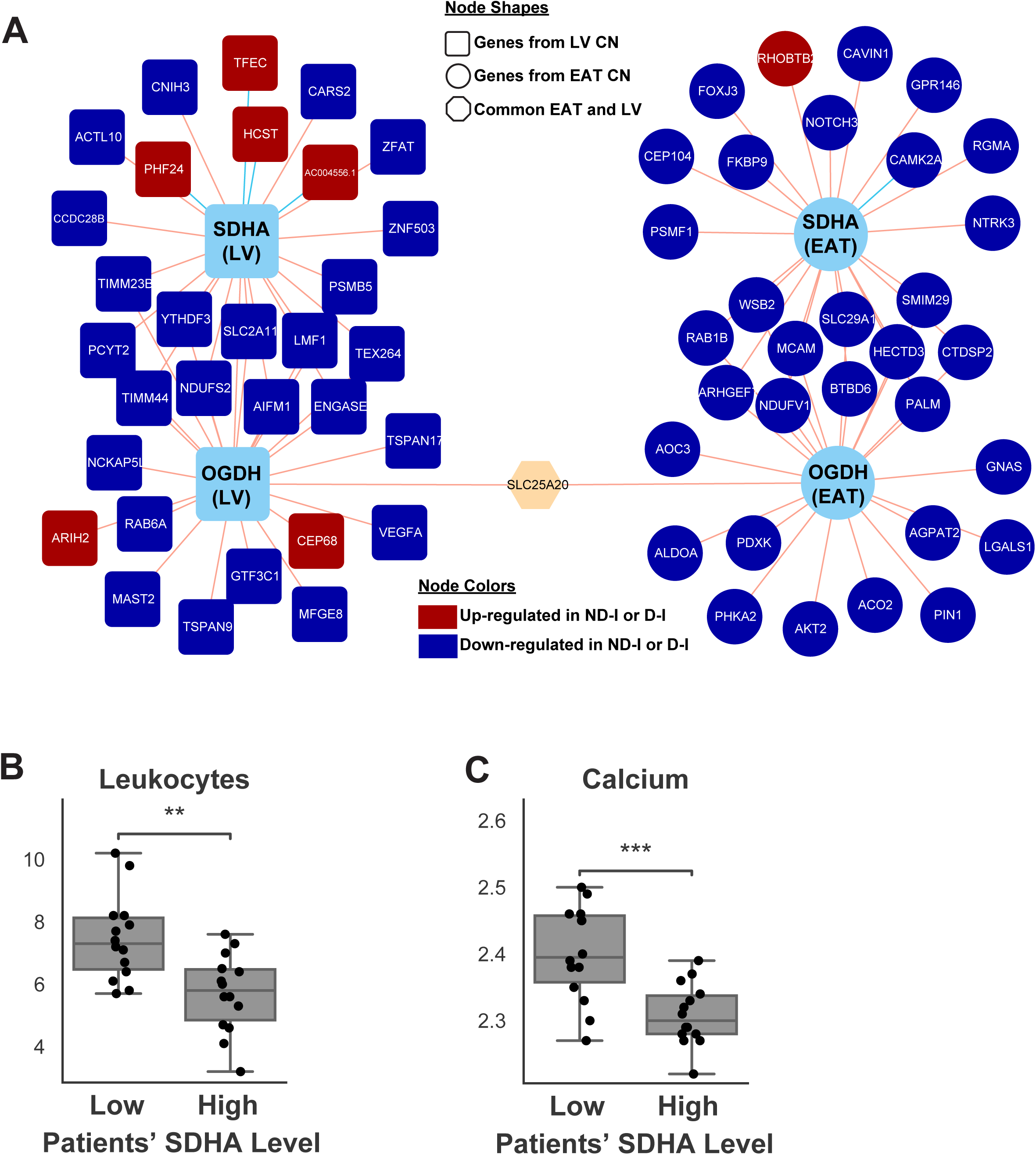
SDHA as a candidate therapeutic target for ischemic heart disease. **(A)** The direct neighbors of SDHA from LV and EAT-specific CN. (B) Leukocyte and Calcium level changes in subjects with low SDHA compared to the subjects with high SDHA in LV

To further understand the systemic effect of SDHA, we re-stratified LV and EAT transcriptomics samples to low and high SDHA (bottom and top 1/3 or 33.3 percentile, respectively) and compared the important clinical variables (**Data S1)** between low and high SDHA groups. We found that the subjects with low LV SDHA levels have significantly higher leukocyte and calcium (**Figure 5B-C**). High levels of these clinical variables are known to be associated with higher inflammations, metabolic dysfunctions, and cardiovascular problems. These correlations further emphasized the importance of the proposed genes to cardiovascular and physiological health.

### Low cardiac SDHA and OGDH level are associated with adverse outcomes of ischemic heart disease

To test our hypothesis on the central role of SDHA and OGDH in the LV and EAT-specific CNs and their association with important clinical variables, we used publicly available heart LV data from GTEx. We selected subjects older than 60 years old, to replicate the population from our study (**Table 1**) and stratified them to low and high SDHA and OGDH (bottom and top 1/3 or 33.3 percentile of TPM values, respectively, for both genes) with 41 samples in high and 47 in the low group (GTEx ID of the selected samples can be found in **Data S6**). The high group consisted of 27 male and 14 female subjects with an average SDHA TPM value of 432.35 ± 11.13 (mean ± SEM) and OGDH of 311.07 ± 10.69 (**Figure 6A**). Meanwhile, the low group consisted of 33 male and 14 female subjects with an average SDHA TPM value of 38.39 ± 3.69 and OGDH of 27.29 ± 2.29 (**Figure 6A**). We performed differential expression and functional analysis between the low and high groups and found that many pathways that were altered in ND-I and D-I subjects compared to healthy were worsened in the low SDHA and OGDH group (**Figure 6B**). We observed the down-regulation of metabolic pathways, including glycolysis, fructose and mannose metabolism, lysine degradation, fatty acid degradation, and TCA cycle, and other important pathways, including oxidative phosphorylation and peroxisome (**Figure 6B**). Moreover, similar trends were followed by many mitochondrial-related biological processes (**Data S6**). Meanwhile, pathways and biological processes related to inflammatory response were up-regulated, including cytokine-cytokine receptor interaction, TNF, NF-kappa B, chemokine, JAK-stat, and B-cell receptor signaling pathways, natural killer cell-mediated toxicity, and Fc gamma R-mediated phagocytosis (**Figure 6B, Data S6**).

**Figure 6:**
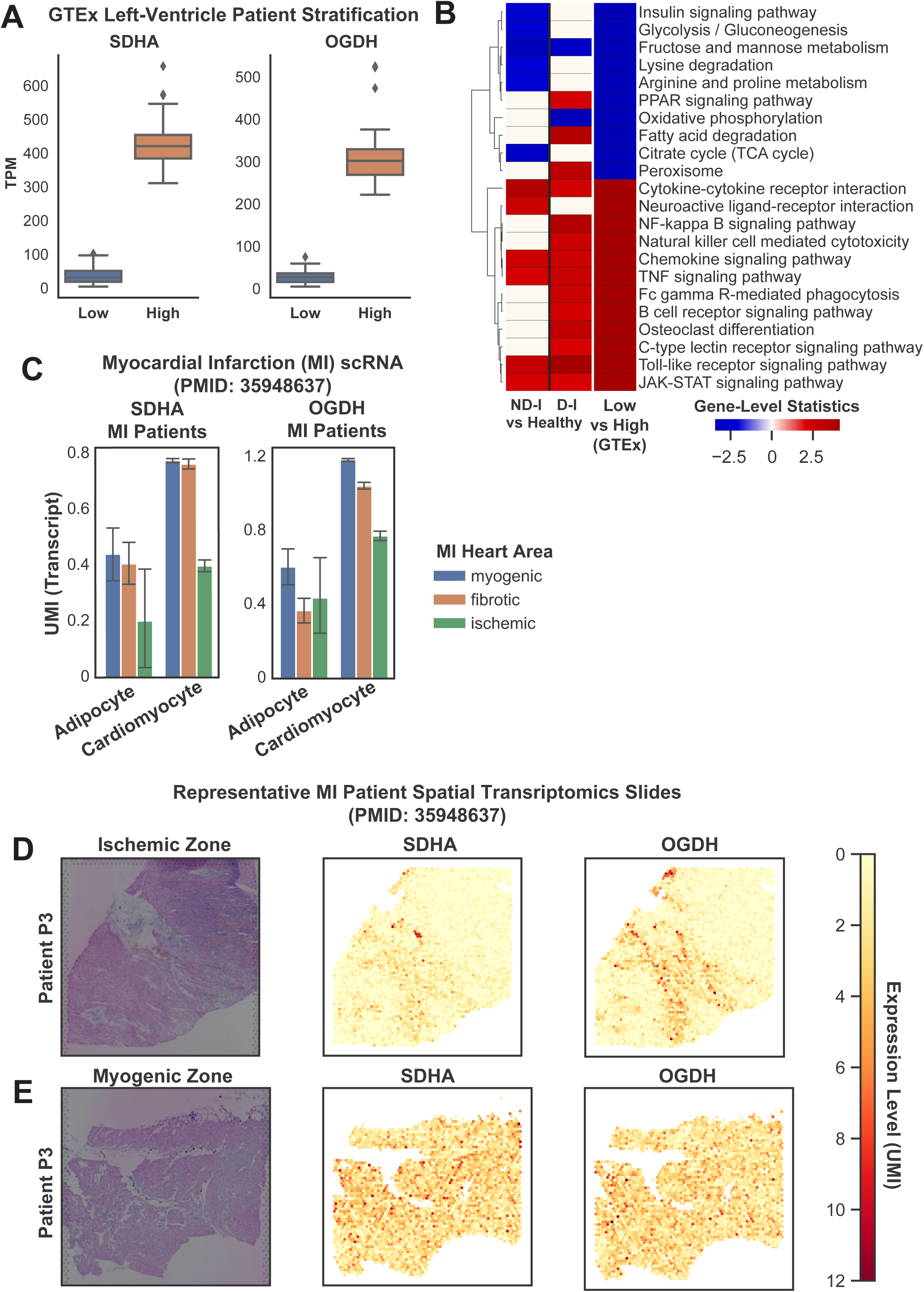
SDHA and OGDH in publicly available heart LV data. **(A)** TPM of low and high SDHA and OGDH levels from heart LV GTEx data. **(B)** Functional analysis of subjects with high SDHA and OGDH compared to the low group. **(C)** The UMI expression of SDHA and OGDH in the single-cell transcriptomics data of MI patients in the two highest expressing cell types: adipocytes and cardiomyocytes. **(D)** Representative spatial transcriptomics expression of SDHA and OGDH from an MI patient (P3) from ischemic and **(E)** myogenic zone.

### Single-cell and spatial transcriptomics data show SDHA and OGDH cell-specificity and emphasizes their involvement in the pathophysiology of ischemic heart disease

By highlighting the central role of SDHA and OGDH in the LV- and EAT-specific CNs, and their association with important clinical variables and pathways in IHD, our analysis demonstrated the strength of network-based approaches to capture meaningful biological signal in heterogeneous patient population, that might be missed by differential expression analysis. This has also been demonstrated in previous studies (Arif *et al*, 2025; Poirel *et al*, 2013). In line with this, we observed modest down-regulations in both LV and EAT of ND-I and D-I compared to healthy (P > 0.05), with a similar trend in our validation cohorts. We hypothesized that this attenuation is due to sample heterogeneity (e.g. disease severity, lifestyle, dietary intake, medications), as we observed significant down-regulations of SDHA and OGDH in bulk transcriptomics data in three controlled mouse studies (Arif *et al*., 2021a; Ounzain *et al*., 2015; Williams *et al*, 2021). To further test our hypothesis, we retrieved publicly available human heart single-cell data that included healthy and myocardial infarction (MI) subjects (Kuppe *et al*, 2022).

We observed that SDHA was expressed (Unique Molecular Identifier or UMI > 0) mainly in cardiomyocytes (70.1% of healthy and 60.4% of MI) and then followed by adipocytes (26.7% in healthy and 33.1% in MI) (**Figure S6**). Focusing on those cell types, we observed similar small down-regulation in SDHA levels as in bulk transcriptomics data, around 16.4% decrease in cardiomyocyte in MI compared to healthy (0.736 ± 0.003, mean ± SEM of UMI per cell vs 0.881 ± 0.005), and 10% decrease in adipocytes (0.397 ± 0.029 vs 0.441 ± 0.199) (**Figure S6**). But when we investigated deeper in ischemic compared to the non-ischemic (myogenic) heart zone of the MI patients, we observed much bigger down-regulation in the SDHA level in almost all cell types (**Figure 6C**).

Specifically, we observed a 48% decrease in cardiomyocytes (0.400 ± 0.011 vs 0.776 ± 0.003 for ischemic and myogenic, respectively) and 54% in adipocytes (0.442 ± 0.091 vs 0.203 ± 0.047). Interestingly, the SDHA level in other necrotic areas of the heart, specifically the fibrotic area, was only decreasing by 1.6% in cardiomyocytes (0.764 ± 0.009) and 7.9% in adipocytes (0.407 ± 0.038) (**Figure 6C**). Similar trends were observed with the expression of OGDH, where we observed a 34% decrease in cardiomyocytes (0.775 ± 0.013 vs 1.187 ± 0.004 for ischemic and myogenic, respectively) and 27% in adipocytes (0.400 ± 0.108 vs 0.607 ± 0.050) (**Figure 6C**). Moreover, in the fibrotic area, the expression of OGDH decreased by 11% in cardiomyocytes (1.046 ± 0.009) and 39% in adipocytes (0.369 ± 0.034) (**Figure 6C**). These observations were validated by spatial transcriptomics data from one of the patients (P3) from the same cohort, where we observed lower SDHA and OGDH expression in the ischemic area (**Figure 6D**) compared to the myogenic area (**Figure 6E**).

## Discussion

Ischemic heart disease is a complex and systemic disease, affecting millions of people in the world. In this study, we generated transcriptomics and employed an extensive systems and network biology analysis to generate a comprehensive transcriptomics profile of ischemic heart disease. One unique feature of our study was that we collected not only heart left ventricle (LV), but also matching epicardial adipose tissue (EAT) from age and BMI-concordant healthy and ischemic subjects. Another unique feature is the inclusion of well-characterized diabetic and non-diabetic ischemic patients based on their clinical records and serum chemistry (i.e. glucose level) data. Subsequently, we generated transcriptomics data and performed differential expression and functional analysis to identify heart-tissue subtype-specific transcriptional and biological changes associated with the disease, including inflammatory and metabolic pathways.

Furthermore, we leveraged co-expression network (CN) analysis to identify driver clusters of each tissue subtype and explore their shared regulatory patterns. Through these analyses, we identified SDHA and OGDH as key genes and potential therapeutic targets and biomarkers for ischemic heart diseases. To emphasize the relevance of SDHA and OGDH to the disease, we stratified heart transcriptomics data from GTEx to high and low SDHA and OGDH groups and compared them. Finally, we validated our proposed targets in previously published mouse and human single-cell and spatial transcriptomics data.

When comparing healthy and ischemic subjects, we observed EAT exhibited more pronounced transcriptional changes compared to LV in response to the disease. This increase is particularly notable in diabetic patients. This observation implies that EAT could serve as a more effective tissue subtype for assessing cardiovascular health and monitoring ischemia progression, particularly in diabetic patients, consistent with established literature (Christensen *et al*, 2020; Iacobellis, 2022; Li *et al*, 2019). Nonetheless, we found many similar affected pathways and functions between the two tissue subtypes, especially for diabetic subjects. Our functional analyses, consistent with the literature (Christodoulidis *et al*, 2014; Koenig, 2001; Mehta & Li, 1999), revealed significant up-regulation of inflammatory pathways, such as cytokine-cytokine receptor interaction, Toll-like receptor, chemokine, NF-kappaB, and TNF signaling, in both regions. Similarly, we also observed down-regulation in oxidative phosphorylation and metabolic pathways, specifically amino acid and carbohydrate metabolism pathways, in both regions. Interestingly, we detected an opposite regulation of the fatty acid degradation pathway in the LV and EAT, which suggests an interplay between the regions as noted in the literature (Fillmore *et al*, 2014; Kankaanpaa *et al*, 2006).

Our co-expression network (CN) analyses provided additional emphasized these shared characteristics and interplay between the two regions. We showed that the driving forces behind the LV and EAT networks consisted of genes associated with crucial biological pathways including oxidative phosphorylation, TCA cycle, lipid metabolism, amino acid metabolism, and fibrosis-related pathways, i.e. focal adhesion and ECM-receptor interaction. These pathways have been studied extensively due to their significance in the pathogenesis of ischemic heart disease (Ferrari, 1996; Kania *et al*, 2009; Liedtke, 1981; Rosca *et al*, 2008). Additionally, according to reporter metabolite analysis, the central network clusters were both enriched with genes linked to NAD and ubiquinol-related metabolites. Notably, reporter transcription factor (TF) analysis indicated that these central clusters also featured target genes of established cardiac TFs.

Another significant result of our CN analysis was the discovery of two promising candidate biomarkers and therapeutic targets: SDHA (succinate dehydrogenase complex flavoprotein subunit A) and OGDH (oxoglutarate dehydrogenase). Both genes were situated in the central clusters (LV-2 and EAT-3), linked to critical ischemia-related functions and pathways, and regulated by known cardiac TFs (SRF and TBX5). Moreover, they are heart-enhanced based on the Human Protein Atlas. Furthermore, we validated their expression down-regulations in multiple different independent cohorts in mice and humans (Arif *et al*., 2021a; Kuppe *et al*., 2022; Ounzain *et al*., 2015; Williams *et al*., 2018) using bulk, single-cell, and spatial transcriptomics data. Additionally, it is worth noting that both genes are FDA-approved and safe druggable targets.

SDHA is a key component of mitochondrial complex II that is essential in the electron transport chain, energy generation, and TCA cycle. Specifically, it is involved in the conversion of succinate to fumarate, FAD to FADH2, and the transfer of electrons from succinate to ubiquinone (Renkema *et al*, 2015). SDHA has been implicated in multiple diseases, including different cancer types (Dubard Gault *et al*, 2018), mitochondrial disorders (Fullerton *et al*, 2020), and Leigh syndrome (Horvath *et al*, 2006). SDHA mutation has also been linked to genetic heart diseases (Courage *et al*., 2017; Levitas *et al*., 2010) and dilated cardiomyopathy (Wang *et al*, 2022). Moreover, using our serum and plasma chemistry data, we observed that subjects with low SDHA have significantly higher leukocyte and calcium compared to the high group. High leukocyte has been used as a signal for internal inflammation in the body and its increase has been linked to the severity of cardiovascular diseases (CVD) (Barron *et al*, 2000; Kim *et al*, 2017; Lee *et al*, 2001). Similarly, the use of calcium supplementation, which implies elevating the level of calcium in the body, is also associated with a higher risk of cardiovascular mortality (Bolland *et al*, 2010; Michaelsson *et al*, 2013).

OGDH functions as a rate-limiting enzyme in the TCA cycle, specifically in the conversion of 2-oxoglutarate to succinyl coenzyme A, and tryptophan metabolism. Recent publications showed a significant decrease in cardiac OGDH levels in subjects with hypertrophic cardiomyopathy (Ranjbarvaziri *et al*., 2021). An obesity-related pre-clinical study showed that the OGDH+/-mice have higher body and liver weight, and heightened susceptibility to liver dysfunction when subjected to high-fat diets (Fan *et al*, 2018). Other publications showed the increase of OGDH and citrate synthase levels in the adrenal gland were proposed as potential biomarkers for atherosclerosis under chronic stress (Meng *et al*, 2021) and the involvement of OGDH in several cancer, developmental, and neurological disorders (Dunckelmann *et al*, 2000; Guffon *et al*, 1993; Sullivan *et al*, 2013; Yap *et al*, 2021).

However, the roles of SDHA and OGDH in ischemic heart disease are still relatively unknown. To illuminate their potential roles, we stratified the heart transcriptomics data from GTEx data to high and low SDHA and OGDH groups. Our analysis unveiled notable patterns: within the low SDHA and OGDH groups, pathways associated with inflammation exhibited up-regulation in the low group compared to the high. These pathways include cytokine-cytokine receptor interaction, NF-kappaB, chemokine signaling, TNF signaling, Toll-like receptor signaling, and JAK-STAT signaling pathways. Conversely, metabolic pathways were down-regulated in the low group, including lysine and fatty acid degradation, TCA cycle, fructose and mannose metabolism, and glycolysis/glucogenesis. In summary, this analysis underscored the exacerbation of vital pathways impacted by ischemic heart disease due to low SDHA and OGDH expression. This suggests a potential role for SDHA and OGDH as potential therapeutics aimed at alleviating the effects of ischemic heart disease.

Although our study offers valuable insights, it is important to acknowledge certain limitations. First, our analysis heavily depended on transcriptomics data, potentially constraining the depth of our findings’ sensitivity. Additionally, our analyses focused on protein-coding genes, overlooking the substantial impact of ischemic heart disease on non-protein coding genes (Busch *et al*, 2016; Poller *et al*, 2018). This study also relied on the correlational relationship between the genes and the disease, not causal relationships. Future investigations hold considerable potential to enhance this study’s scope by encompassing other omics (e.g. metabolomics, proteomics, and metagenomics) in both bulk and single cell level and non-protein coding genes in their exploration.

In summary, our study provides valuable insights into the molecular landscape of ischemic heart disease and unveils promising candidates for further investigation and development of therapeutic interventions. We employed systems biology and data-driven approaches to systematically study the molecular, specifically transcriptional, alterations and dysregulation caused by the disease, in both diabetic and non-diabetic subjects. We also used co-expression network analysis to identify central clusters and genes associated with the disease. Finally, we identified SDHA and OGDH as potential biomarkers and therapeutic targets to attenuate the damage caused by the disease, highlighting their crucial roles in the pathogenesis of ischemic heart disease.

## STAR Methods

### KEY RESOURCES TABLE

| Deposited Data |  |  |
| --- | --- | --- |
| RNA-Seq Data | this paper | Data S1 |
| Clinical and Demography Data | this paper | Data S1 |
| RNA-Seq Data | PMID: 33972017 | GSE153485 |
| RNA-Seq Data | PMID: 29652636 | GSE104187 |
| RNA-Seq Data | PMID: 24786300 | GSE52313 |
| Single-Cell and Spatial Transcriptomics | PMID: 35948637 | https://zenodo.org/record/6578047 |
| Software and Algorithms |  |  |
| Python 3.7 | Python Package | RRID:SCR_019053 |
| R | R Project for Statistical Computing | RRID:SCR_001905 |
| Kallisto | PMID:27043002 | RRID:SCR_016582 |
| Rpy2 | Python Package | https://rpy2.github.io/ |
| DESeq2 | R Package | RRID:SCR_015687 |
| PIANO | R Package | RRID:SCR_003200 |
| SciPy | Python Package | RRID:SCR_008058 |
| Statsmodel | Python Package | RRID:SCR_016074 |
| iGraph | Python Package | RRID:SCR_019225 |
| Leiden Clustering | Python Package | https://github.com/vtraag/leidenalg |
| Statsmodel | Python Package | RRID:SCR_016074 |
| Gseapy | Python Package | https://gseapy.readthedocs.io/en/latest/ |
| Enrichr | Python Package | RRID:SCR_001575 |

### RESOURCE AVAILABILITY

#### Lead Contact

Further information and requests for resources and reagents should be directed to and will be fulfilled by the lead contact, Muhammad Arif.

#### Materials Availability

This study did not generate new unique reagents; the reagents used are available commercially.

#### Data and Code Availability

The processed transcriptomics data (raw counts and TPM), clinical, and demography data can be retrieved from **Data S1**. The code for the analysis and visualization (grouped based on the figure numbers) is available at https://github.com/muharif/2025.Arif_Doran_etal_IschemicHeartDisease.

### EXPERIMENTAL MODEL AND SUBJECT DETAILS

#### Sample Collections from Human Subjects

Cardiac ischemic biopsies were obtained from 30 patients undergoing coronary artery bypass surgery due to significant atherosclerotic blockage in the epicardial coronary arteries. All patients were examined through echocardiography prior to operation and the ejection fraction (EF) was measured. Non-ischemic biopsies from the left ventricle were obtained from 14 subjects undergoing aortic valve replacement, with angiography verified absence of coronary artery disease in any of the major myocardial coronary artery branches. All biopsies were collected from the left ventricle septum region with a 1 mm needle. The average weight of collected tissues samples was 100 mg and these were stored at -80°C until analysis. All patients gave informed and written consent. The study was approved by the Gothenburg Regional Ethics Committee and done according to the Declaration of Helsinki.

### METHOD DETAILS

#### Clinical and demography data analysis

The clinical and demography data were compared using one-way ANOVA (*“f_oneway”*) with Tukey’s post-hoc (*“pairwise_tukeyhsd”*) test. All of these analysis modules were provided by the *Statsmodel* (Seabold & Perktold, 2010) v0.13.2. We used the *“ttest_ind”* function when comparing these variables in the low and high SDHA and OGDH groups (**Figure 5B-C**) from SciPy (Virtanen *et al*, 2020) package version 1.7.3. Variables with q-values < 0.05 were considered significant in both analyses.

#### RNA extraction and sequencing

Total RNA was isolated from biopsies relating to 44 samples from the left ventricle using RNeasy Fibrous Tissue Mini kit (QIAGEN) and 24 samples from epicardial fat using RNeasy Lipid Tissue Mini kit (QIAGEN). These 68 samples were processed with SMARTer® Stranded RNA-Seq Kit (Takara Bio) for reverse transcription, generation of double stranded cDNA and subsequent library preparation. All libraries were quantified with the Fragment Analyzer using the standard sensitivity NGS kit (Agilent Technologies), pooled in equimolar concentrations and quantified with a Qubit Fluorometer (ThermoFisher Scientific), the library pool was further diluted and sequenced using 150 cycles on an Illumina NextSeq500.

#### Transcriptomics Data Analysis

The paired-end RNA-sequencing raw files were quantified using Kallisto (Bray *et al*, 2016) with an index file generated from the Ensembl human reference genome (Cunningham *et al*, 2022) and mapped to gene-level count and TPM using the mapping from Ensembl Biomart website, by selecting only the protein-coding transcripts and genes. Differential expression and functional analysis were performed using an in-house *DESeq2* (Love *et al*, 2014) and Piano (Varemo *et al*, 2013) wrapper in Python, using the *rpy2* version 3.2.2 package. The multiple hypothesis testing was done within the packages with Benjamini-Hochburg correction. We used *seaborn* (Waskom et al, 2017) version 0.11.2 package to visualize the results. The KEGG and Gene Ontology biological processes gene set collections were retrieved from *Enrichr* (Chen *et al*, 2013; Kuleshov *et al*, 2016), specifically the “Human 2021” version.

#### Co-Expression Network Analysis

Co-expression networks were generated using Spearman correlation rank, from *SciPy* version 1.7.3 package (Virtanen *et al*., 2020), in each tissue subtype using the TPM count data, after removing the bottom 25% of the least varying genes. Gene-gene correlation with (1) Benjamin-Hochberg FDR < 0.05 and (2) among the top 25% positively correlated genes was kept for further analysis (“*multipletests*” function from Statsmodel v0.13.2). Network analyses were done with iGraph (Antonov *et al*, 2023) version 0.9.11 and community detection/clustering analysis was done using Leiden (Traag *et al*., 2019) algorithm in iGraph. *Enrichr* (Chen *et al*., 2013; Kuleshov *et al*., 2016) was used for the clusters’ functional analysis.

#### Single-Cell and Spatial Transcriptomics Visualization

The single-cell and spatial transcriptomics data were retrieved from their respective data repository and the pre-processed h5ad files were loaded using *scanpy* (Wolf *et al*, 2018) version 1.9.3 package, without any further processing. The UMI bar plots were generated using the *Seaborn* version 0.11.2 package.

## Supplementary Information

**Figure S1:**
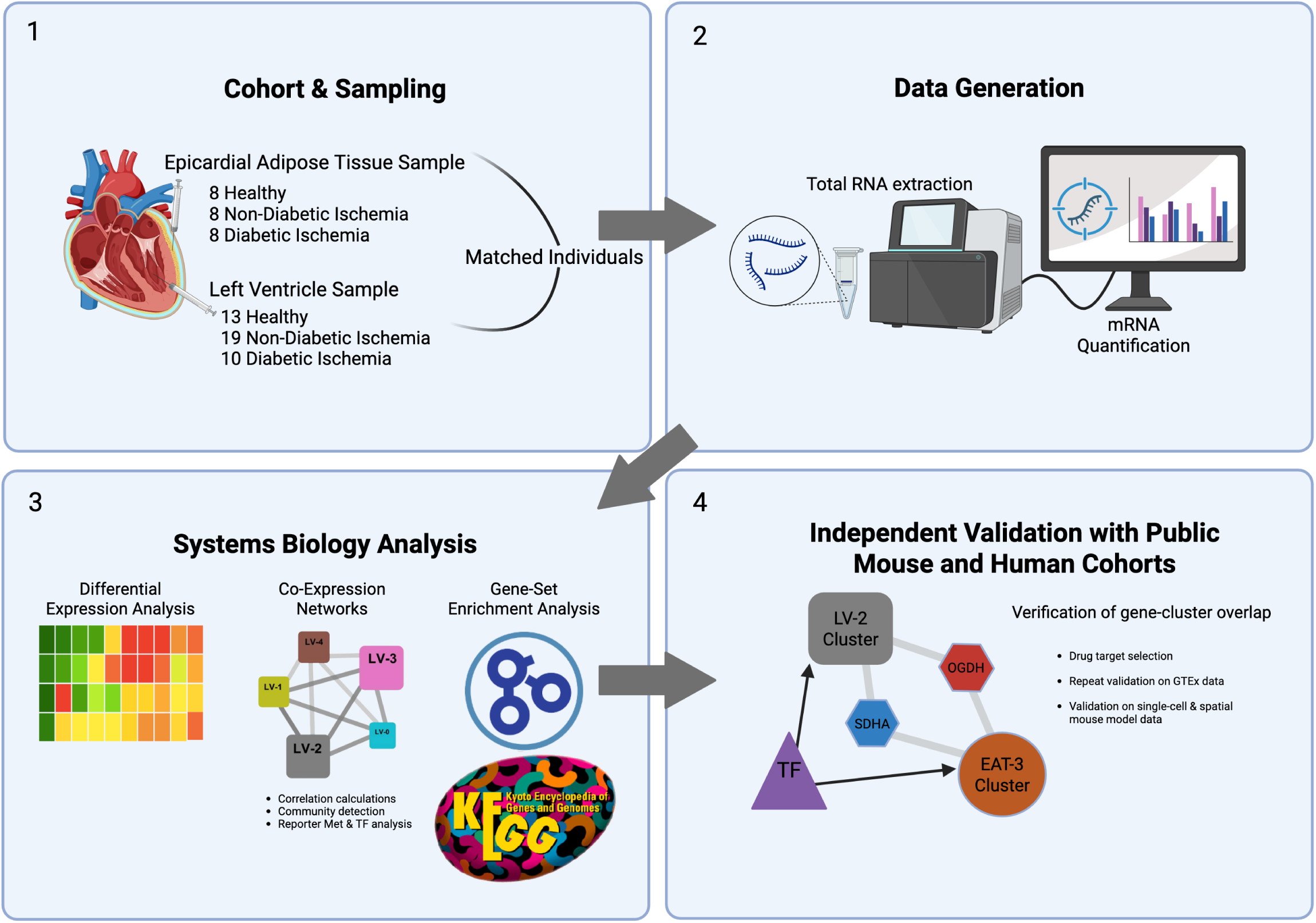
Graphical Abstract and Study Flow

**Figure S2:**
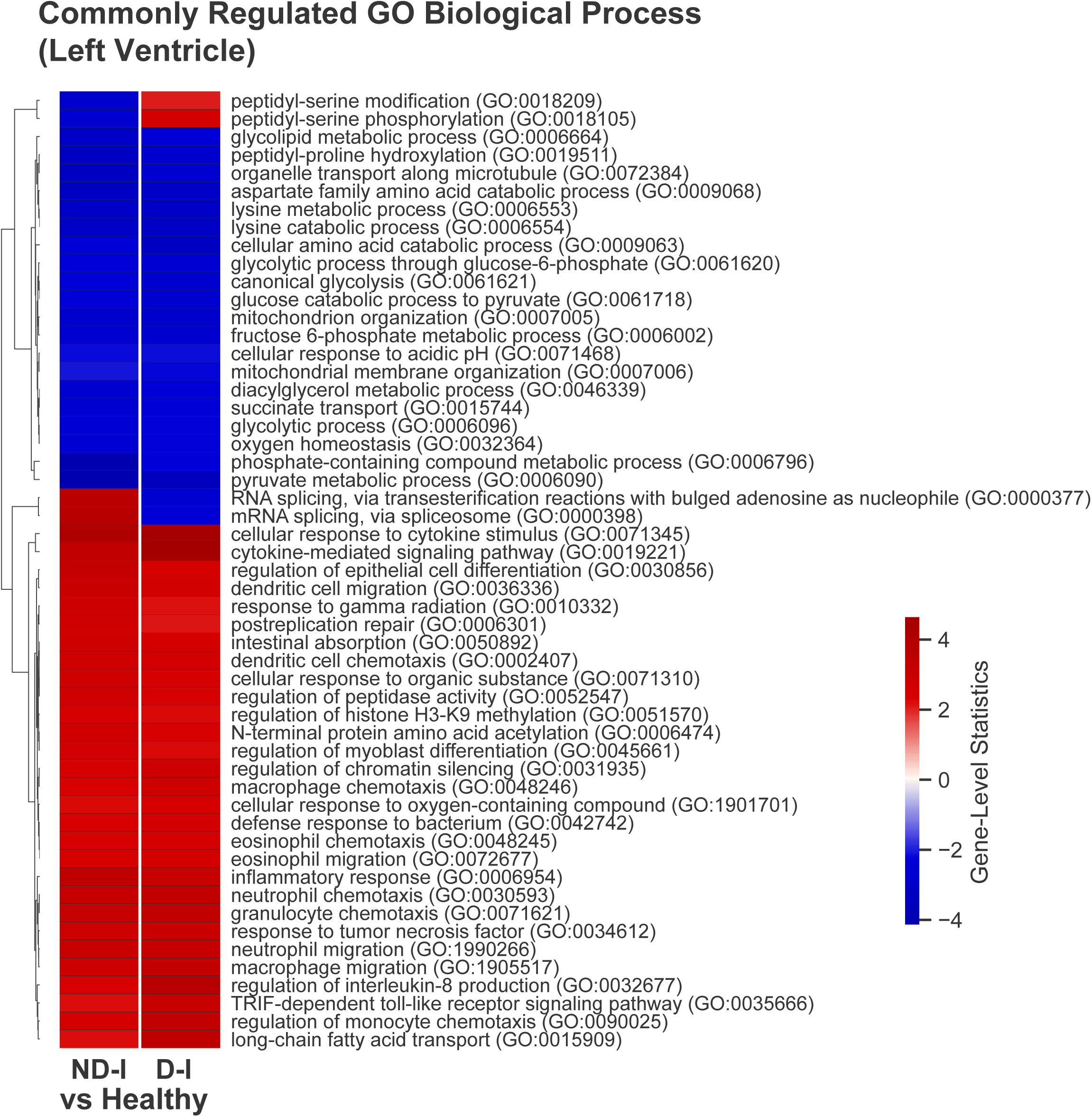
Commonly altered gene ontology biological process (P-Value < 0.01) between ischemic (ND-I and D-I) and healthy groups from Heart Left Ventricle transcriptomics data (Related to Figure 1).

**Figure S3:**
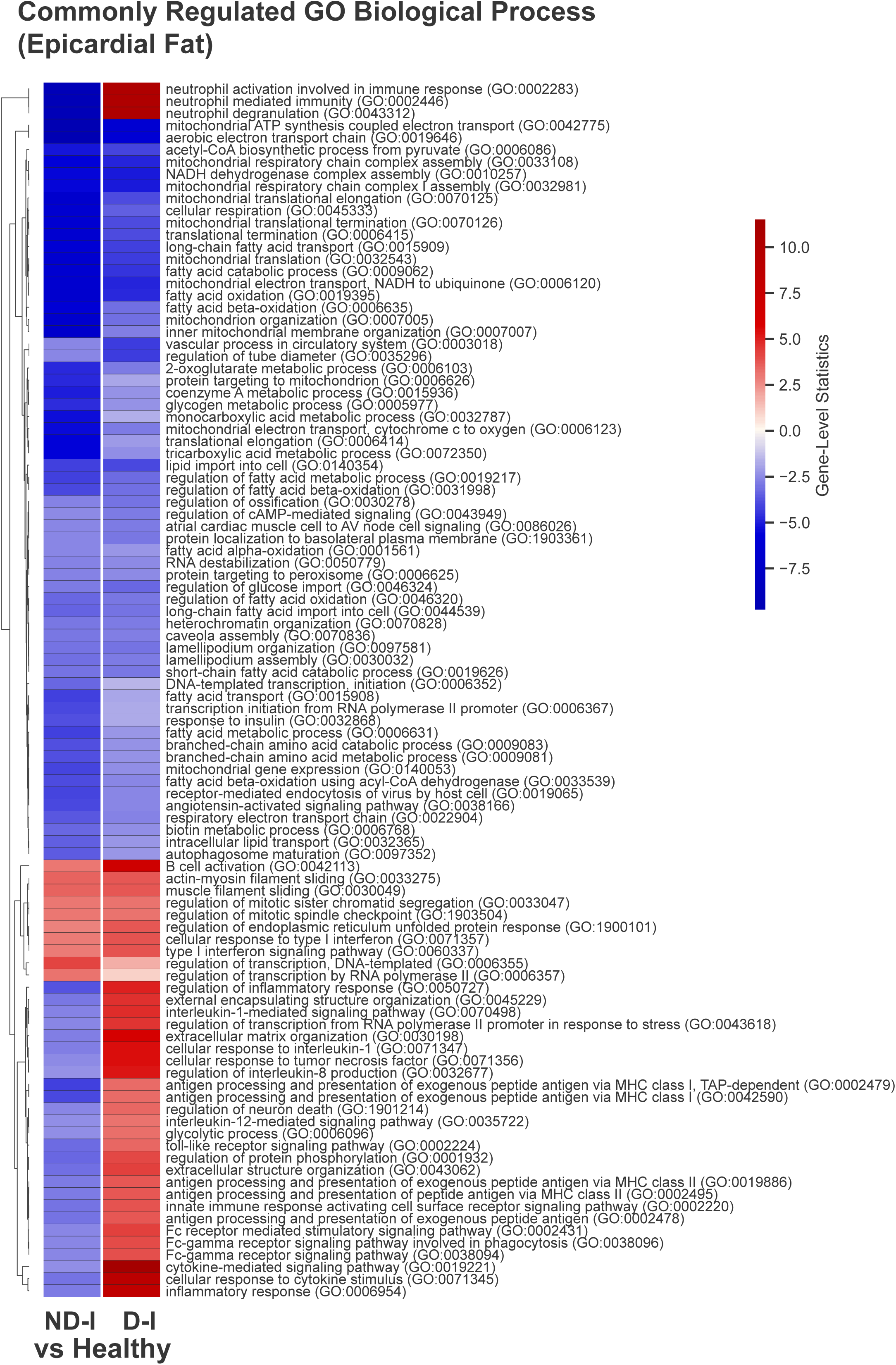
Commonly altered gene ontology biological process (P-Value < 0.01) between ischemic (ND-I and D-I) and healthy groups from Epicardial Adipose TIssue transcriptomics data (Related to Figure 2).

**Figure S4:**
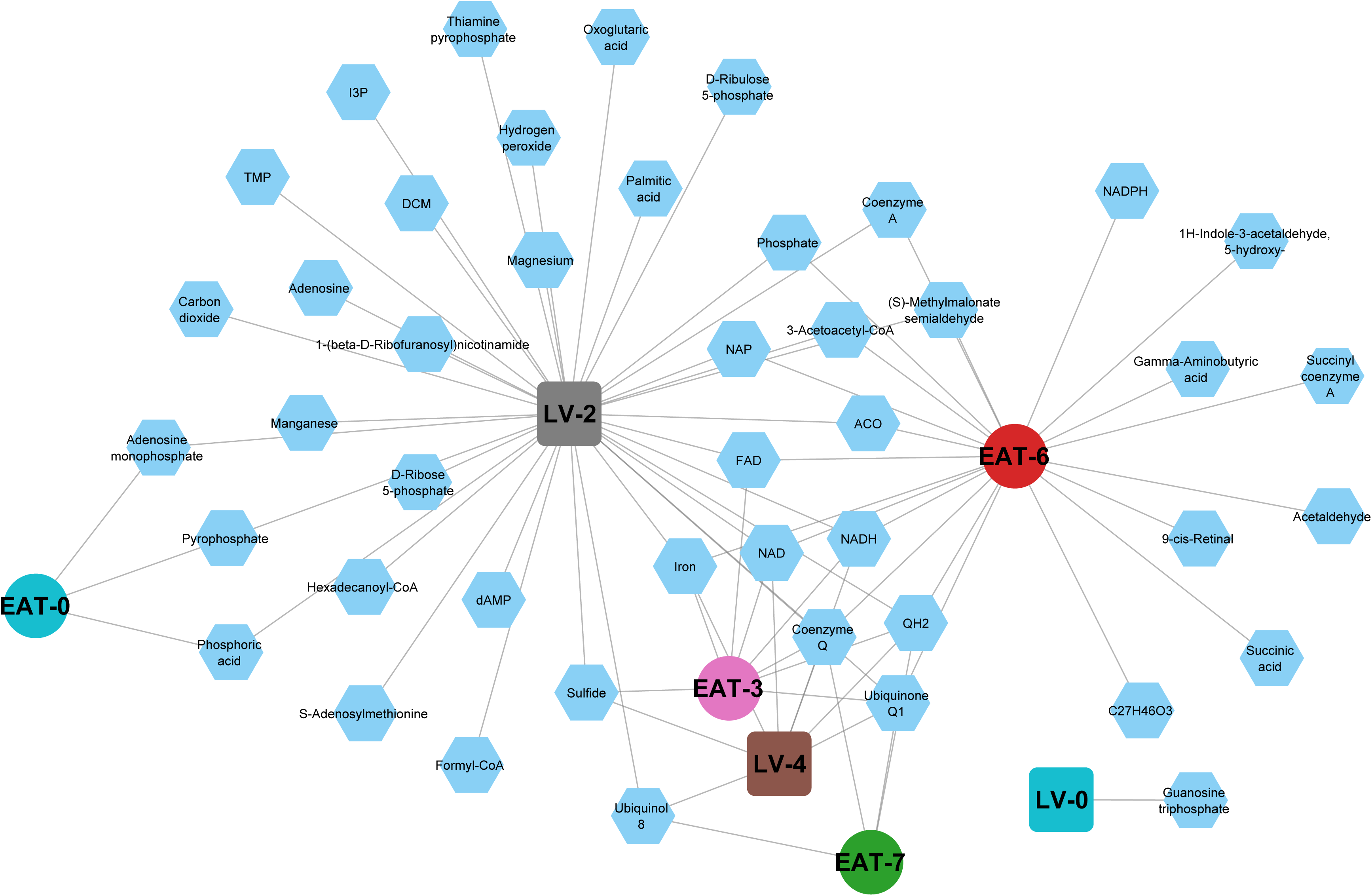
Reporter metabolite analysis results connected to their associated CN clusters. For visualization purposes, we removed 763 diacylglycerol subtypes and 21 unnamed metabolites that were connected to LV-2 (Related to Figure 4).

**Figure S5:**
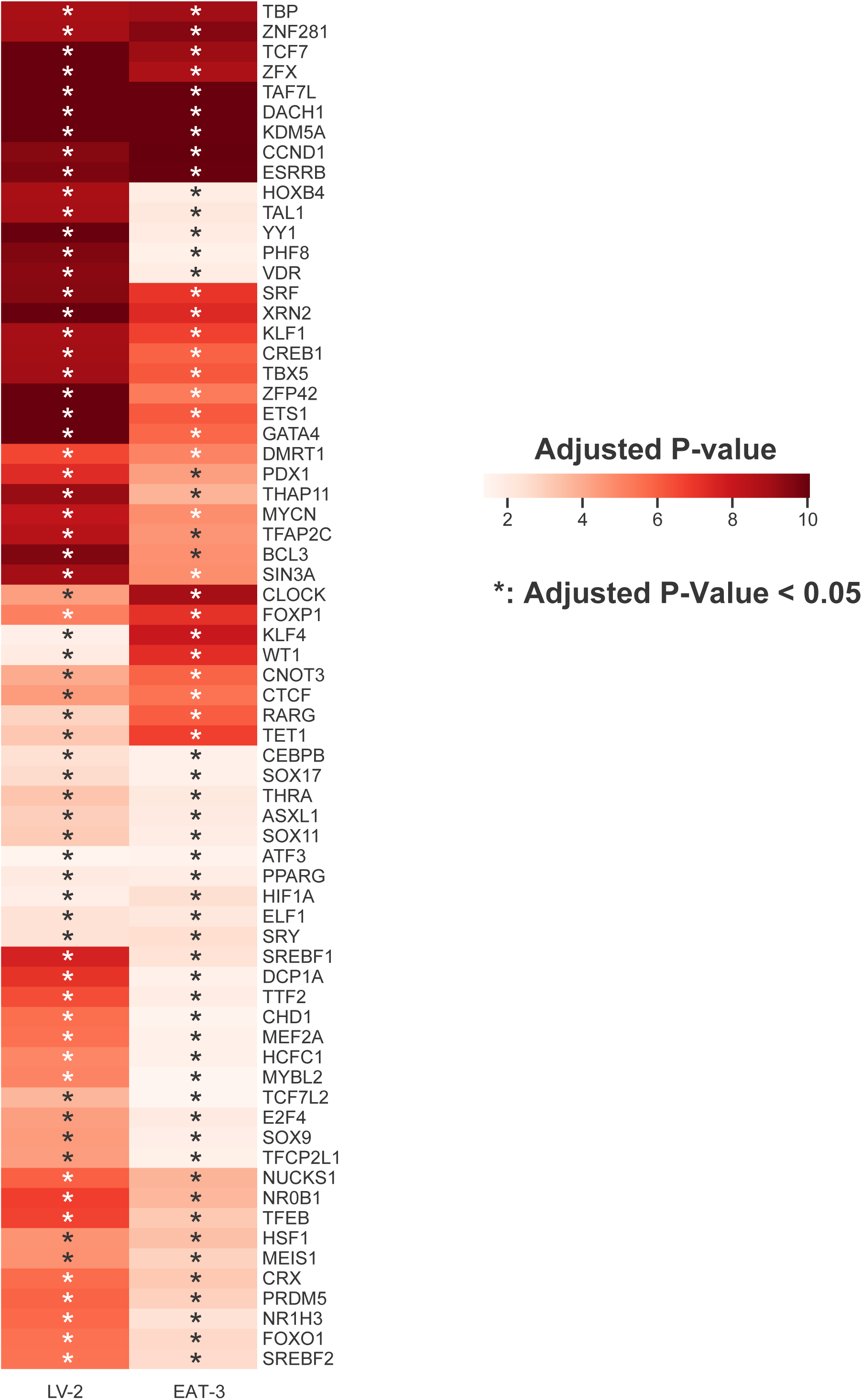
Common reporter transcription factor (TF) results between LV-2 and EAT-3. The annotations denoted the number of known TF target genes from the clusters (Related to Figure 4).

**Figure S6:**
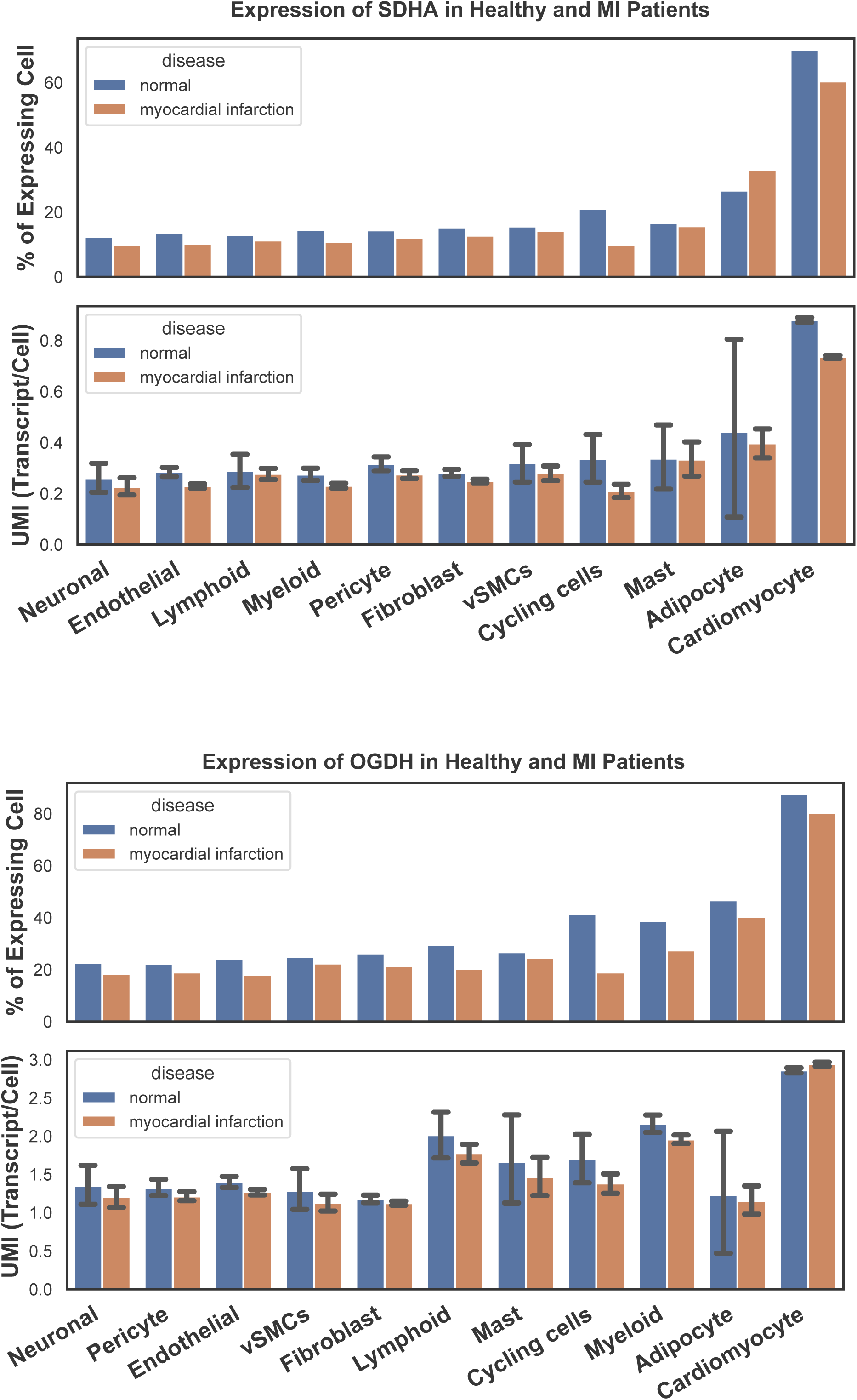
Single-cell expression of SDHA and OGDH when comparing in healthy and myocardial infarction heart (Related to Figure 6).

## Acknowledgements

This work was supported by the SciLifeLab & Wallenberg Data Driven Life Science Program (M.A., grant: KAW 2020.0239). The computations and data handling were enabled by resources provided by the National Academic Infrastructure for Supercomputing in Sweden (NAISS), partially funded by the Swedish Research Council through grant agreement no. 2022-06725.

## References

Antonov M, Csárdi G, Horvát S, Müller K, Nepusz T, Noom D, Salmon M, Traag V, Welles BF, Zanini F (2023) igraph enables fast and robust network analysis across programming languages. arXiv preprint arXiv:2311 10260

Arif M, Basu A, Wolf KM, Park JK, Pommerolle L, Behee M, Gochuico BR, Cinar R (2023) An Integrative Multiomics Framework for Identification of Therapeutic Targets in Pulmonary Fibrosis. Adv Sci (Weinh) 10: e2207454

Arif M, Klevstig M, Benfeitas R, Doran S, Turkez H, Uhlen M, Clausen M, Wikstrom J, Etal D, Zhang C et al (2021a) Integrative transcriptomic analysis of tissue-specific metabolic crosstalk after myocardial infarction. Elife 10

Arif M, Lehoczki A, Hasko G, Lohoff FW, Ungvari Z, Pacher P (2025) Global and tissue-specific transcriptomic dysregulation in human aging: Pathways and predictive biomarkers. Geroscience 47: 5917–5936

Arif M, Zhang C, Li X, Gungor C, Cakmak B, Arslanturk M, Tebani A, Ozcan B, Subas O, Zhou W et al (2021b) iNetModels 2.0: an interactive visualization and database of multi-omics data. Nucleic Acids Res 49: W271–W276

Barron HV, Cannon CP, Murphy SA, Braunwald E, Gibson CM (2000) Association between white blood cell count, epicardial blood flow, myocardial perfusion, and clinical outcomes in the setting of acute myocardial infarction: a thrombolysis in myocardial infarction 10 substudy. Circulation 102: 2329–2334

Bertero E, Maack C (2018) Metabolic remodelling in heart failure. Nat Rev Cardiol 15: 457–470

Bolland MJ, Avenell A, Baron JA, Grey A, MacLennan GS, Gamble GD, Reid IR (2010) Effect of calcium supplements on risk of myocardial infarction and cardiovascular events: meta-analysis. BMJ 341: c3691

Bray NL, Pimentel H, Melsted P, Pachter L (2016) Near-optimal probabilistic RNA-seq quantification. Nat Biotechnol 34: 525–527

Brown DA, Perry JB, Allen ME, Sabbah HN, Stauffer BL, Shaikh SR, Cleland JG, Colucci WS, Butler J, Voors AA et al (2017) Expert consensus document: Mitochondrial function as a therapeutic target in heart failure. Nat Rev Cardiol 14: 238–250

Busch A, Eken SM, Maegdefessel L (2016) Prospective and therapeutic screening value of non-coding RNA as biomarkers in cardiovascular disease. Ann Transl Med 4: 236

CDC, 2024. Heart Disease Facts.

Chen EY, Tan CM, Kou Y, Duan Q, Wang Z, Meirelles GV, Clark NR, Ma’ayan A (2013) Enrichr: interactive and collaborative HTML5 gene list enrichment analysis tool. BMC Bioinformatics 14: 128

Christensen RH, von Scholten BJ, Lehrskov LL, Rossing P, Jorgensen PG (2020) Epicardial adipose tissue: an emerging biomarker of cardiovascular complications in type 2 diabetes? Ther Adv Endocrinol Metab 11: 2042018820928824

Christodoulidis G, Vittorio TJ, Fudim M, Lerakis S, Kosmas CE (2014) Inflammation in coronary artery disease. Cardiol Rev 22: 279–288

Courage C, Jackson CB, Hahn D, Euro L, Nuoffer JM, Gallati S, Schaller A (2017) SDHA mutation with dominant transmission results in complex II deficiency with ocular, cardiac, and neurologic involvement. Am J Med Genet A 173: 225–230

Cunningham F, Allen JE, Allen J, Alvarez-Jarreta J, Amode MR, Armean IM, Austine-Orimoloye O, Azov AG, Barnes I, Bennett R et al (2022) Ensembl 2022. Nucleic Acids Res 50: D988–D995

Dubard Gault M, Mandelker D, DeLair D, Stewart CR, Kemel Y, Sheehan MR, Siegel B, Kennedy J, Marcell V, Arnold A et al (2018) Germline SDHA mutations in children and adults with cancer. Cold Spring Harb Mol Case Stud 4

Dunckelmann RJ, Ebinger F, Schulze A, Wanders RJ, Rating D, Mayatepek E (2000) 2-ketoglutarate dehydrogenase deficiency with intermittent 2-ketoglutaric aciduria. Neuropediatrics 31: 35–38

Fan Z, Li L, Li X, Zhang M, Zhong Y, Li Y, Yu D, Cao J, Zhao J, Xiaoming D et al (2018) Generation of an oxoglutarate dehydrogenase knockout rat model and the effect of a high-fat diet. RSC Adv 8: 16636–16644

Ferrari R (1996) The role of mitochondria in ischemic heart disease. J Cardiovasc Pharmacol 28 Suppl 1: S1–10

Fillmore N, Mori J, Lopaschuk GD (2014) Mitochondrial fatty acid oxidation alterations in heart failure, ischaemic heart disease and diabetic cardiomyopathy. Br J Pharmacol 171: 2080–2090

Fullerton M, McFarland R, Taylor RW, Alston CL (2020) The genetic basis of isolated mitochondrial complex II deficiency. Mol Genet Metab 131: 53–65

Guffon N, Lopez-Mediavilla C, Dumoulin R, Mousson B, Godinot C, Carrier H, Collombet JM, Divry P, Mathieu M, Guibaud P (1993) 2-Ketoglutarate dehydrogenase deficiency, a rare cause of primary hyperlactataemia: report of a new case. J Inherit Metab Dis 16: 821–830

Horvath R, Abicht A, Holinski-Feder E, Laner A, Gempel K, Prokisch H, Lochmuller H, Klopstock T, Jaksch M (2006) Leigh syndrome caused by mutations in the flavoprotein (Fp) subunit of succinate dehydrogenase (SDHA). J Neurol Neurosurg Psychiatry 77: 74–76

Huang M, Bao J, Hallstrom BM, Petranovic D, Nielsen J (2017) Efficient protein production by yeast requires global tuning of metabolism. Nat Commun 8: 1131

Iacobellis G (2022) Epicardial adipose tissue in contemporary cardiology. Nat Rev Cardiol 19: 593–606

Jiang DS, Zeng HL, Li R, Huo B, Su YS, Fang J, Yang Q, Liu LG, Hu M, Cheng C et al (2017) Aberrant Epicardial Adipose Tissue Extracellular Matrix Remodeling in Patients with Severe Ischemic Cardiomyopathy: Insight from Comparative Quantitative Proteomics. Sci Rep 7: 43787

Kania G, Blyszczuk P, Eriksson U (2009) Mechanisms of cardiac fibrosis in inflammatory heart disease. Trends Cardiovasc Med 19: 247–252

Kankaanpaa M, Lehto HR, Parkka JP, Komu M, Viljanen A, Ferrannini E, Knuuti J, Nuutila P, Parkkola R, Iozzo P (2006) Myocardial triglyceride content and epicardial fat mass in human obesity: relationship to left ventricular function and serum free fatty acid levels. J Clin Endocrinol Metab 91: 4689–4695

Kim JH, Lim S, Park KS, Jang HC, Choi SH (2017) Total and differential WBC counts are related with coronary artery atherosclerosis and increase the risk for cardiovascular disease in Koreans. PLoS One 12: e0180332

Koenig W (2001) Inflammation and coronary heart disease: an overview. Cardiol Rev 9: 31–35

Kuleshov MV, Jones MR, Rouillard AD, Fernandez NF, Duan Q, Wang Z, Koplev S, Jenkins SL, Jagodnik KM, Lachmann A et al (2016) Enrichr: a comprehensive gene set enrichment analysis web server 2016 update. Nucleic Acids Res 44: W90–97

Kuppe C, Ramirez Flores RO, Li Z, Hayat S, Levinson RT, Liao X, Hannani MT, Tanevski J, Wunnemann F, Nagai JS et al (2022) Spatial multi-omic map of human myocardial infarction. Nature 608: 766–777

Lee CD, Folsom AR, Nieto FJ, Chambless LE, Shahar E, Wolfe DA (2001) White blood cell count and incidence of coronary heart disease and ischemic stroke and mortality from cardiovascular disease in African-American and White men and women: atherosclerosis risk in communities study. Am J Epidemiol 154: 758–764

Lee S, Zhang C, Liu Z, Klevstig M, Mukhopadhyay B, Bergentall M, Cinar R, Stahlman M, Sikanic N, Park JK et al (2017) Network analyses identify liver-specific targets for treating liver diseases. Mol Syst Biol 13: 938

Levitas A, Muhammad E, Harel G, Saada A, Caspi VC, Manor E, Beck JC, Sheffield V, Parvari R (2010) Familial neonatal isolated cardiomyopathy caused by a mutation in the flavoprotein subunit of succinate dehydrogenase. Eur J Hum Genet 18: 1160–1165

Li Y, Liu B, Li Y, Jing X, Deng S, Yan Y, She Q (2019) Epicardial fat tissue in patients with diabetes mellitus: a systematic review and meta-analysis. Cardiovasc Diabetol 18: 3

Liedtke AJ (1981) Alterations of carbohydrate and lipid metabolism in the acutely ischemic heart. Prog Cardiovasc Dis 23: 321–336

Liu M, Lv J, Pan Z, Wang D, Zhao L, Guo X (2022) Mitochondrial dysfunction in heart failure and its therapeutic implications. Front Cardiovasc Med 9: 945142

Liu X, Yuan M, Zhao D, Zeng Q, Li W, Li T, Li Q, Zhuo Y, Luo M, Chen P et al (2024) Single-Nucleus Transcriptomic Atlas of Human Pericoronary Epicardial Adipose Tissue in Normal and Pathological Conditions. Arterioscler Thromb Vasc Biol 44: 1628–1645

Love MI, Huber W, Anders S (2014) Moderated estimation of fold change and dispersion for RNA-seq data with DESeq2. Genome Biol 15: 550

Mahabadi AA, Berg MH, Lehmann N, Kalsch H, Bauer M, Kara K, Dragano N, Moebus S, Jockel KH, Erbel R et al (2013) Association of epicardial fat with cardiovascular risk factors and incident myocardial infarction in the general population: the Heinz Nixdorf Recall Study. J Am Coll Cardiol 61: 1388–1395

Mahabadi AA, Lehmann N, Kalsch H, Robens T, Bauer M, Dykun I, Budde T, Moebus S, Jockel KH, Erbel R et al (2014) Association of epicardial adipose tissue with progression of coronary artery calcification is more pronounced in the early phase of atherosclerosis: results from the Heinz Nixdorf recall study. JACC Cardiovasc Imaging 7: 909–916

Mehta JL, Li DY (1999) Inflammation in ischemic heart disease: response to tissue injury or a pathogenetic villain? Cardiovasc Res 43: 291–299

Meng LB, Hu GF, Shan MJ, Zhang YM, Yu ZM, Liu YQ, Xu HX, Wang L, Gong T, Liu DP (2021) Citrate Synthase and OGDH as Potential Biomarkers of Atherosclerosis under Chronic Stress. Oxid Med Cell Longev 2021: 9957908

Michaelsson K, Melhus H, Warensjo Lemming E, Wolk A, Byberg L (2013) Long term calcium intake and rates of all cause and cardiovascular mortality: community based prospective longitudinal cohort study. BMJ 346: f228

Mohar DS, Salcedo J, Hoang KC, Kumar S, Saremi F, Erande AS, Naderi N, Nadeswaran P, Le C, Malik S (2014) Epicardial adipose tissue volume as a marker of coronary artery disease severity in patients with diabetes independent of coronary artery calcium: findings from the CTRAD study. Diabetes Res Clin Pract 106: 228–235

Moskowitz DP (1986) Reform of the state professional disciplinary process: five modest proposals. N Y State Dent J 52: 32–34

Oikonomou EK, Marwan M, Desai MY, Mancio J, Alashi A, Hutt Centeno E, Thomas S, Herdman L, Kotanidis CP, Thomas KE et al (2018) Non-invasive detection of coronary inflammation using computed tomography and prediction of residual cardiovascular risk (the CRISP CT study): a post-hoc analysis of prospective outcome data. Lancet 392: 929–939

Ounzain S, Micheletti R, Beckmann T, Schroen B, Alexanian M, Pezzuto I, Crippa S, Nemir M, Sarre A, Johnson R et al (2015) Genome-wide profiling of the cardiac transcriptome after myocardial infarction identifies novel heart-specific long non-coding RNAs. Eur Heart J 36: 353–368a

Poirel CL, Rahman A, Rodrigues RR, Krishnan A, Addesa JR, Murali TM (2013) Reconciling differential gene expression data with molecular interaction networks. Bioinformatics 29: 622–629

Poller W, Dimmeler S, Heymans S, Zeller T, Haas J, Karakas M, Leistner DM, Jakob P, Nakagawa S, Blankenberg S et al (2018) Non-coding RNAs in cardiovascular diseases: diagnostic and therapeutic perspectives. Eur Heart J 39: 2704–2716

Ranjbarvaziri S, Kooiker KB, Ellenberger M, Fajardo G, Zhao M, Vander Roest AS, Woldeyes RA, Koyano TT, Fong R, Ma N et al (2021) Altered Cardiac Energetics and Mitochondrial Dysfunction in Hypertrophic Cardiomyopathy. Circulation 144: 1714–1731

Renkema GH, Wortmann SB, Smeets RJ, Venselaar H, Antoine M, Visser G, Ben-Omran T, van den Heuvel LP, Timmers HJ, Smeitink JA et al (2015) SDHA mutations causing a multisystem mitochondrial disease: novel mutations and genetic overlap with hereditary tumors. Eur J Hum Genet 23: 202–209

Rosca MG, Vazquez EJ, Kerner J, Parland W, Chandler MP, Stanley W, Sabbah HN, Hoppel CL (2008) Cardiac mitochondria in heart failure: decrease in respirasomes and oxidative phosphorylation. Cardiovasc Res 80: 30–39

Seabold S, Perktold J, 2010. statsmodels: Econometric and statistical modeling with python, 9th Python in Science Conference.

Sullivan LB, Martinez-Garcia E, Nguyen H, Mullen AR, Dufour E, Sudarshan S, Licht JD, Deberardinis RJ, Chandel NS (2013) The proto-oncometabolite fumarate binds glutathione to amplify ROS-dependent signaling. Mol Cell 51: 236–248

Traag VA, Waltman L, van Eck NJ (2019) From Louvain to Leiden: guaranteeing well-connected communities. Sci Rep 9: 5233

Uhlen M, Fagerberg L, Hallstrom BM, Lindskog C, Oksvold P, Mardinoglu A, Sivertsson A, Kampf C, Sjostedt E, Asplund A et al (2015) Proteomics. Tissue-based map of the human proteome. Science 347: 1260419

Varemo L, Nielsen J, Nookaew I (2013) Enriching the gene set analysis of genome-wide data by incorporating directionality of gene expression and combining statistical hypotheses and methods. Nucleic Acids Res 41: 4378–4391

Virtanen P, Gommers R, Oliphant TE, Haberland M, Reddy T, Cournapeau D, Burovski E, Peterson P, Weckesser W, Bright J et al (2020) SciPy 1.0: Fundamental Algorithms for Scientific Computing in Python. Nature Methods 17: 261–272

Wang CP, Hsu HL, Hung WC, Yu TH, Chen YH, Chiu CA, Lu LF, Chung FM, Shin SJ, Lee YJ (2009) Increased epicardial adipose tissue (EAT) volume in type 2 diabetes mellitus and association with metabolic syndrome and severity of coronary atherosclerosis. Clin Endocrinol (Oxf) 70: 876–882

Wang X, Zhang X, Cao K, Zeng M, Fu X, Zheng A, Zhang F, Gao F, Zou X, Li H et al (2022) Cardiac disruption of SDHAF4-mediated mitochondrial complex II assembly promotes dilated cardiomyopathy. Nat Commun 13: 3947

Waskom M, Botvinnik O, O’Kane D, Hobson P, Lukauskas S, Gemperline DC, Augspurger T, Halchenko Y, Cole JB, Warmenhoven J et al, 2017. mwaskom/seaborn: v0.8.1 (September 2017), v0.8.1 ed. Zenodo.

WHO, 2025. Cardiovascular diseases (CVDs).

Williams AL, Khadka V, Tang M, Avelar A, Schunke KJ, Menor M, Shohet RV (2018) HIF1 mediates a switch in pyruvate kinase isoforms after myocardial infarction. Physiol Genomics 50: 479–494

Williams AL, Walton CB, Pinell B, Khadka VS, Dunn B, Lee K, Anagaran MCT, Avelar A, Shohet RV (2021) Ischemic heart injury leads to HIF1-dependent differential splicing of CaMK2gamma. Sci Rep 11: 13116

Wolf FA, Angerer P, Theis FJ (2018) SCANPY: large-scale single-cell gene expression data analysis. Genome Biol 19: 15

Yap ZY, Strucinska K, Matsuzaki S, Lee S, Si Y, Humphries K, Tarnopolsky MA, Yoon WH (2021) A biallelic pathogenic variant in the OGDH gene results in a neurological disorder with features of a mitochondrial disease. J Inherit Metab Dis 44: 388–400

Zeybel M, Arif M, Li X, Altay O, Yang H, Shi M, Akyildiz M, Saglam B, Gonenli MG, Yigit B et al (2022) Multiomics Analysis Reveals the Impact of Microbiota on Host Metabolism in Hepatic Steatosis. Adv Sci (Weinh) 9: e2104373

Zhou B, Tian R (2018) Mitochondrial dysfunction in pathophysiology of heart failure. J Clin Invest 128: 3716–3726

Zou R, Zhang M, Zou Z, Shi W, Tan S, Wang C, Xu W, Jin J, Milton S, Chen Y et al (2023) Single-cell transcriptomics reveals zinc and copper ions homeostasis in epicardial adipose tissue of heart failure. Int J Biol Sci 19: 4036–4051

